# Evolutionarily conserved effects of Notch signaling drive intestinal graft-versus-host disease in mice and non-human primates

**DOI:** 10.1101/2022.04.23.488844

**Authors:** Victor Tkachev, Ashley Vanderbeck, Eric Perkey, Scott N. Furlan, Connor McGuckin, Daniela Gómez Atria, Ulrike Gerdemann, Xianliang Rui, Jennifer Lane, Daniel J. Hunt, Hengqi Zheng, Lucrezia Colonna, Michelle Hoffman, Alison Yu, Samantha Kelly, Anneka Allman, Brandon Burbach, Yoji Shimizu, Angela Panoskaltsis-Mortari, Guoying Chen, Stephen M. Carpenter, Olivier Harari, Frank Kuhnert, Gavin Thurston, Bruce R. Blazar, Leslie S. Kean, Ivan Maillard

## Abstract

Notch signaling promotes T-cell pathogenicity and graft-versus-host disease (GVHD) after allogeneic hematopoietic cell transplantation (allo-HCT) in mice, with a dominant role for the Delta-like ligand DLL4. To assess if Notch’s effects are evolutionarily conserved and identify key mechanisms, we studied antibody-mediated DLL4 blockade in a non-human primate model similar to human allo-HCT. Short-term DLL4 blockade improved post-transplant survival with striking, durable protection from gastrointestinal GVHD, out of proportion to other disease sites. Unlike prior immunosuppressive strategies, anti-DLL4 interfered with a T-cell transcriptional program associated with intestinal infiltration. In cross-species investigations, Notch inhibition decreased surface abundance of the gut-homing integrin a4b7 in conventional T-cells via b1 competition for a4 binding, while preserving a4b7 in regulatory T-cells. Thereby, DLL4/Notch blockade decreased effector T-cell infiltration into the gut, with increased regulatory to conventional T-cell ratios early after allo-HCT. Our results identify a conserved, biologically unique and targetable role of DLL4/Notch signaling in GVHD.

**One Sentence Summary:** Notch signaling promotes pathogenic effector T cell infiltration of the intestine during acute graft-versus-host disease.

## INTRODUCTION

Allogeneic hematopoietic cell transplantation (allo-HCT) has life-saving potential for patients with hematological malignancies and bone marrow disorders. However, acute graft-versus-host disease (aGVHD) remains a major cause of morbidity and mortality (*1*). Among aGVHD manifestations, gastrointestinal involvement (GI-aGVHD) is one of the most challenging, with nearly all cases of severe aGVHD prominently involving the gastrointestinal tract. Moreover, epithelial injury and the gut microbiome can fuel activation of pathogenic T cells, thereby propagating tissue damage after allo-HCT (*2–8*). Thus, preventing GI-aGVHD would represent a major advance if accomplished while preserving protective immunity, including anti-infectious and anti-cancer T cell responses (*9*).

To foster progress, preclinical studies have relied heavily on mouse allo-HCT models. Major advantages include strain combinations that mimic many aspects of aGVHD, as well as abundant genetic and immunological reagents (*10*). Yet, mouse models do not account for evolutionary changes in the importance of biological pathways driving aGVHD, especially for complex ligand/receptor systems with redundant family members. Moreover, mouse allo-HCT differs from human allo-HCT in terms of transplant conditioning, supportive care and complications. Thus, multiple factors could underlie the failure of preclinical studies in mice to be translated to patients. To address these issues, we developed a T cell-replete haploidentical allo-HCT model of aGVHD in non-human primates (NHP) (*11–16*). In this model, new treatments can be evaluated initially as single agents for their activity in GVHD prophylaxis, unlike in human patients where new interventions must be combined with drugs used for routine GVHD prevention. Thus, the impact of targeting a single pathway can be studied in detail in NHP (*12, 13, 15, 16*), with key questions for subsequent translation: a) What is the magnitude of single agent activity?, b) Is the pathway of interest conserved from mice to primates, and worth targeting in humans?, c) Are there unexpected toxicities not predicted in mice?, and d) Can we identify unique mechanisms of GVHD protection informing how new strategies should be deployed in humans? We previously reported that early post-transplant Notch inhibition in donor T cells induced major protection from GVHD in mice (*17–21*). Notch is an evolutionarily conserved ligand/receptor signaling pathway. Mammals harbor four agonistic Notch ligands (Delta-like1/4, Jagged1/2) and four receptors (Notch1-4), with both redundant and non-redundant functions (*22, 23*). Ligand/receptor interactions lead to proteolytic cleavage of the Notch receptor by g- secretase, followed by translocation of intracellular Notch to the nucleus, where it induces target gene activation in association with the transcription factor RBP-Jk and a Mastermind-like family co-activator. In mouse allo-HCT models, genetic or pharmacologic Notch inhibition in donor T cells led to substantial protection from aGVHD, while preserving T cell expansion in lymphoid organs as well as potent graft-versus-tumor activity – a beneficial pattern of immunomodulation (*17-21, 24-26*). Notch1/2 in T cells and Delta-like1/4 (DLL1/4) in the host accounted for all of the effects of Notch signaling on GVHD, with a dominant role for Notch1 and DLL4 (*17, 21,*

*24*). Donor T cells interacted with Delta-like ligands expressed by specialized subsets of fibroblastic reticular cells in secondary lymphoid organs within hours of transplantation, consistent with an early window of pathogenic Notch activity (*24*). Indeed, Notch inhibition within two days of allo-HCT was essential to confer GVHD protection, and a single dose of antibodies blocking Delta-like ligands provided long-term GVHD protection when given at the time of transplantation (*24*). Thus, our findings in mice suggest that transient inhibition of Delta-like Notch ligands is an attractive new strategy to prevent GVHD after allo-HCT, without causing global immunosuppression.

Despite these promising findings, it remains unknown if the critical impact of Notch signaling on GVHD in mice will be conserved in humans. Although its overall effects are preserved between species, Notch is a complex pathway driven by multiple receptors and ligands whose individual involvement can drift during evolution. Furthermore, systemic Notch inhibitors such as gDsecretase inhibitors induced significant on-target toxicity in mice and in human clinical trials, especially when given continuously (*21, 27–29*). In contrast, targeting individual Notch ligands and receptors with blocking antibodies conferred a broader therapeutic window (*21, 30*). Thus, it is essential to define the overall activity, tolerability, temporal effects and mechanisms-of-action of specific Notch pathway inhibitors in models of allo-HCT and GVHD as close to human allo-HCT as possible.

To address these questions, we turned to our haploidentical allo-HCT model in NHPs, which recapitulates key aspects of human allo-HCT, providing a translational platform for investigational drugs and mechanistic analysis. Using this approach, we nominated new therapeutic strategies first studied in mice as candidates for human translation (*11–16*), with the first (CTLA4-Ig) now FDA-approved for GVHD prevention (*31*). Given the dominant role of the Notch ligand DLL4 in mice, we prioritized transient peri-transplant DLL4 inhibition as the most promising strategy. We tested an anti-DLL4 antibody (REGN421) cross-reactive with human and NHP DLL4 for which pharmacokinetic and toxicity information was available from prior studies in NHP and cancer patients (*32*). In the NHP allo-HCT model, transient DLL4 inhibition with one dose or three weekly doses of REGN421 had potent single agent activity against clinical and pathologic GI-aGVHD, a pattern of activity not observed previously in NHP with other candidate therapeutics. Mechanistic analysis revealed decreased accumulation of activated conventional T cells in the GI tract and an increased ratio of regulatory to conventional T cells, both in NHP and in mice. Notch-deprived effector but not regulatory T cells had decreased surface levels of the gut-homing integrin a4b7, which interacts with endothelial MAdCAM-1 close to intestinal crypts (*5*). Our data suggest a new mechanism for decreased a4b7 levels through enhanced b1 and b7 competition for a4 binding upon Notch blockade. Together, we identified a highly conserved pathogenic role of the Notch ligand DLL4 from mice to NHP, with major therapeutic benefits of transient DLL4 inhibition in preventing GI-aGVHD. Our data provide critical information about the unique effects of Notch inhibition on the differential accumulation of regulatory and conventional T cells in the gut. In turn, these observations provide new insights into the development of Notch targeting strategies to prevent GVHD in patients.

## RESULTS

### DLL4 blockade early after allo-HCT protects NHP from GI-aGVHD

Our prior experiments in mice demonstrated that short-term blockade of Delta-like Notch ligands within 2 days of allo-HCT had substantial protective effects against aGVHD (*21, 24*). To evaluate if these effects translate to NHP, we treated NHP allo-HCT recipients with the anti- DLL4 blocking antibody REGN421, using one of two dosing regimens: a single dose of 3 mg/kg at day 0 post-HCT (n=7), or three weekly doses at day 0, 7 and 14 (n=4). We then compared these NHP to allo-HCT recipients receiving supportive care only (“NoRx”) (Fig. 1A). Pharmacokinetic analysis after single dosing of REGN421 demonstrated mono-phase decay kinetics with a half-life of 3.36 days, resulting in biologically active serum antibody concentrations >2 µg/mL (*33*) until day 17±2 days (Fig. S1A). Three weekly doses of REGN421 also resulted in antibody levels >2 µg/mL for >30 days post-HCT (Fig. S1A).

**Figure 1.**
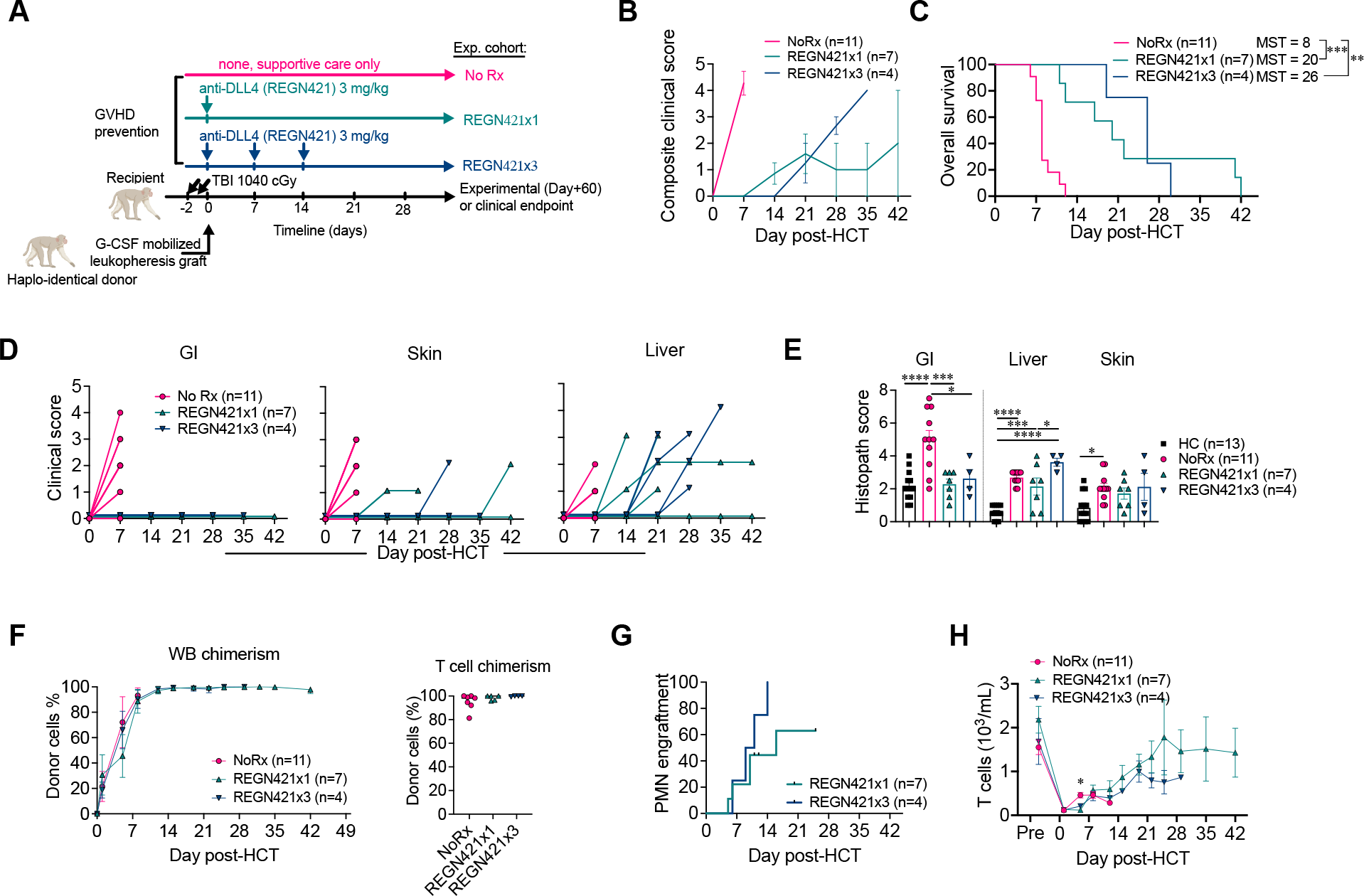
DLL4 blockade early after allo-HCT protects from GI-aGVHD in the NHP model. (A) Experimental design, depicting major components of the NHP aGVHD model and the dosing regimens with a single or three weekly doses of REGN421. (B) Composite clinical score of allo-HCT recipients in NoRx, REGN421x1 and REGN421x3 cohorts. (C) Overall survival of allo-HCT recipients in the NoRx aGVHD cohort (pink, n =11), REGN421x1 (teal, n=7) and REGN421x3 (blue, n=4) experimental cohorts. Recipients euthanized based on pre- determined experimental endpoints were censored at terminal analysis. **p<0.01, ***p<0.001, log-rank (Mantel-Cox) test. (D) Clinical scores for GI, skin and liver aGVHD in allo-HCT recipients, based on established criteria (diarrhea, skin rash and bilirubin level) (*14*). (E) Histopathological aGVHD scores for skin, liver and GI tract (terminal ileum and colon). *p<0.05, ***p<0.001, ****p<0.0001, ANOVA with Tukey post-hoc-test. (F) Donor chimerism in whole blood and CD3^+^/CD20^−^ T cells sorted at terminal analysis. (G) Neutrophil engraftment in REGN421x1 and REGN421x3 cohorts. (H) Absolute number of T cells in the peripheral blood.

Significant protection from aGVHD was observed with both a single dose of REGN421 at Day 0 or three weekly doses at Day 0, 7 and 14, consistent with a potent impact of REGN421 during early phases of T cell activation (Fig. 1B-C). REGN421-treated animals had delayed aGVHD onset and lengthened survival compared to controls with no prophylaxis (NoRx; Fig. 1B-C). Detailed analysis of the aGVHD clinical presentation in REGN421-treated animals in comparison to NoRx controls revealed near complete protection from GI-aGVHD with REGN421. Indeed, none of the eleven recipients that received either single or multiple doses of REGN421 developed clinical signs of GI-aGVHD (Fig. 1D). Moreover, pathology scores in the GI tract were significantly lower with REGN421 compared to those from the NoRx cohort and equivalent to scores in healthy non-transplanted NHP (Fig. 1E). However, REGN421-treated recipients did ultimately develop other clinical and pathologic signs of skin, hepatic and/or pulmonary aGVHD, but without GI disease (Table S1, Fig, 1D-E, Fig. S1B). These observations are consistent with partial protection from aGVHD after short-term DLL4 inhibition, but with strikingly complete protection from GI-aGVHD by clinical and pathological criteria.

Recipients receiving mono-prophylaxis with a single dose or three weekly doses of REGN421 had effective donor engraftment after allo-HCT (Fig. S1D-K), including high levels of bone marrow, whole blood as well as T cell donor chimerism (Fig. 1F, Fig. 1SC). Nine out of eleven recipients showed hematologic reconstitution post-transplant (Fig. 1G-H); two animals (R.306 and R.297, each treated with a single dose of REGN421) were euthanized early after transplant (on Day +12 and +11, respectively, prior to neutrophil engraftment). R.306 exhibited pre-engraftment bleeding complications, and R.297 developed a pre-engraftment GI infection (Table S1), both of which are expected complications after allo-HCT in NHP (*12*) and patients (*34, 35*). Thus, 5 animals in the single-dose cohort and 4 animals in the 3-dose cohort were evaluable for all aGVHD-associated clinical and immunological outcomes. As our 1-dose and 3- dose anti-DLL4 cohorts were similar by clinical, pathological and immunological criteria, all data from these cohorts were merged in subsequent analyses. Together, our findings suggest that DLL4 blockade can be achieved after allo-HCT in NHP without drug-associated toxicity, without interfering with donor engraftment and with high protective activity against GI-aGVHD.

### DLL4 blockade preserves mature T cell differentiation and reconstitution despite preventing GI aGVHD

Our group previously examined OX40L blockade with the KY1005 antibody (*13*) and CD28 blockade with FR104 (*12*) for their ability to prevent T cell activation and prolong aGVHD-free survival in NHP. With these agents, aGVHD prevention correlated with a block in T cell activation and maturation, resulting in substantial relative increases of CD4^+^ and CD8^+^ T cells with a naïve phenotype (Fig. S2A-B). In contrast, flow cytometric analysis showed that, despite substantial protection against GI-aGVHD, REGN421 did not block T cell differentiation after HCT. Indeed, the expected pattern of post-transplant naïve-to-central-memory and effector-memory transitions in CD4^+^ T cells, and naïve-to-effector-memory and terminal-effector transitions in CD8^+^ T cells were similar in animals treated with REGN421 and in the NoRx cohort (Fig. 2A-B, Fig. S2A-B). Although proportions of proliferating Ki67^+^CD4^+^ and CD8^+^ T cells were reduced in REGN421-treated compared to NoRx animals (Fig. 2C), other activation markers were similar between groups. This included similar upregulation of OX40 and PD-1 on blood CD4^+^ T cells (Fig. 2D-E), and upregulation of CD69 on CD8^+^ T cells in both cohorts, with a slightly delayed but preserved upregulation of PD-1 on CD8^+^ T cells in the REGN421 cohort (Fig. 2D-F). Yet, DLL4 inhibition induced similarly prolonged survival after allo-HCT (median survival time [MST] = 22 days) compared with the KY1005 anti-OX40L antibody (MST = 19.5 days) (*13*) and the FR104 anti-CD28 antibody (MST = 21 days) (*12*) (Fig. S2C). Importantly, unlike REGN421, treatment with KY1005 or FR104 did not improve GI pathology compared with the NoRx GVHD cohort (Fig. S2D). Together, these data suggest that Notch inhibition protects allo-HCT recipients from aGVHD by mechanisms other than those of OX40L- or CD28-directed costimulation blockade.

**Figure 2.**
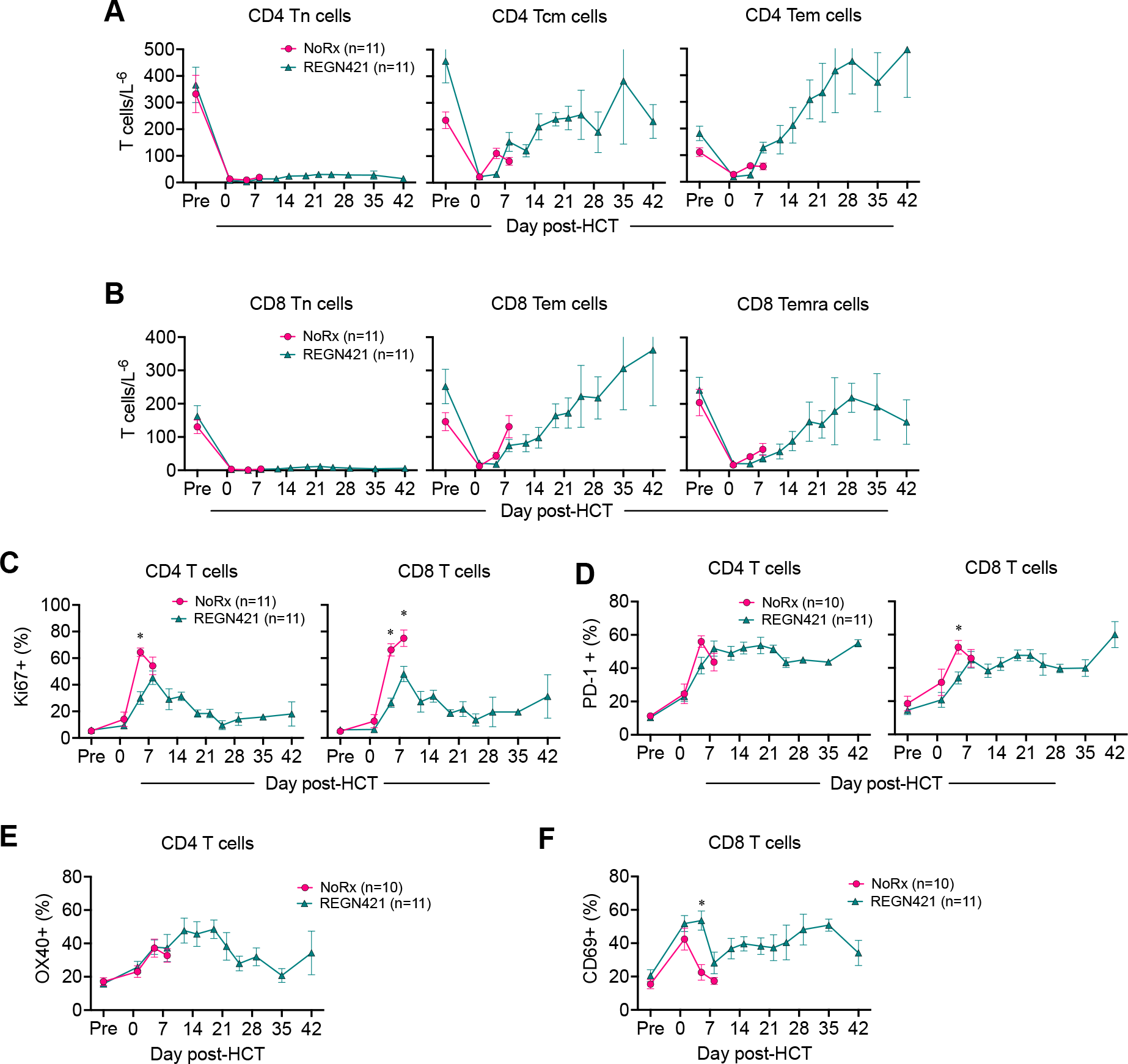
DLL4 blockade does not interfere with T cell differentiation and reconstitution following allo-HCT. (A-B) Absolute number of CD4^+^ (A) and CD8^+^ (B) T cells with CD45RA^+^CCR7^+^CD95^−^ naïve, CD45RA^−^CCR7^+^ central memory, CD45RA^−^CCR7^−^ effector memory and CD45RA^+^CCR7^−^ terminal effector phenotypes in the blood of allo-HCT recipients. *p<0.05, Tukey-corrected t-test. (C-D) Relative number of Ki67^+^ (C) or PD-1^+^ (D) CD4^+^ and CD8^+^ T cells in the blood of allo-HCT recipients. *p<0.05, Tukey-corrected t-test. (E-F) Relative number of OX40^+^CD4^+^ (E) and CD69^+^CD8^+^ (F) T cells in the blood of allo-HCT recipients.

### REGN421 prophylaxis decreases cell surface a4b7 integrin in CD8^+^ T cells after allo-HCT

Given that REGN421 affected T cells in a manner distinct from our previous work, and that it specifically protected from GI but not liver or skin aGVHD, we hypothesized that DLL4 blockade early after allo-HCT altered unique aspects of aGVHD pathogenesis, including migration of activated T cells to the GI tract. To test this hypothesis, we studied homing molecules controlling T cell migration into visceral organs during aGVHD (*36–40*). We analyzed relevant chemokine receptors and integrins using flow cytometry (including CCR5, CCR9 and CXCR3, as well as the integrin a4b7) and quantified mRNA for the chemokine receptor gene *CXCR6* (*37*), for which rhesus macaque-reactive antibodies do not exist. While REGN421 did not impact expression of CCR5, CXCR3, CXCR6 or CCR9 (Fig. S3A-D), it decreased cell surface level of the GI tract-homing integrin a4b7 on CD8^+^ T cells in blood (Fig. 3A-D), spleen (Fig. 3E-F) and mesenteric lymph nodes (mLN; Fig. 3G-H) compared to NoRx controls. Decreased cell surface a4b7 correlated with a lower number of a4b7^+^CD8^+^ T cells also expressing CCR9 in REGN421-treated compared to NoRx controls (Fig. S3E). Importantly, while development of aGVHD in the NoRx cohort was associated with lower a4b7 levels in regulatory T cells (Treg) compared to heathy controls, DLL4 blockade normalized the proportion of a4b7^+^ Treg (Fig. 3G-H). Thus, REGN421 decreased cell surface a4b7 in conventional T cells (Tcon) while preserving it in Treg.

**Figure 3.**
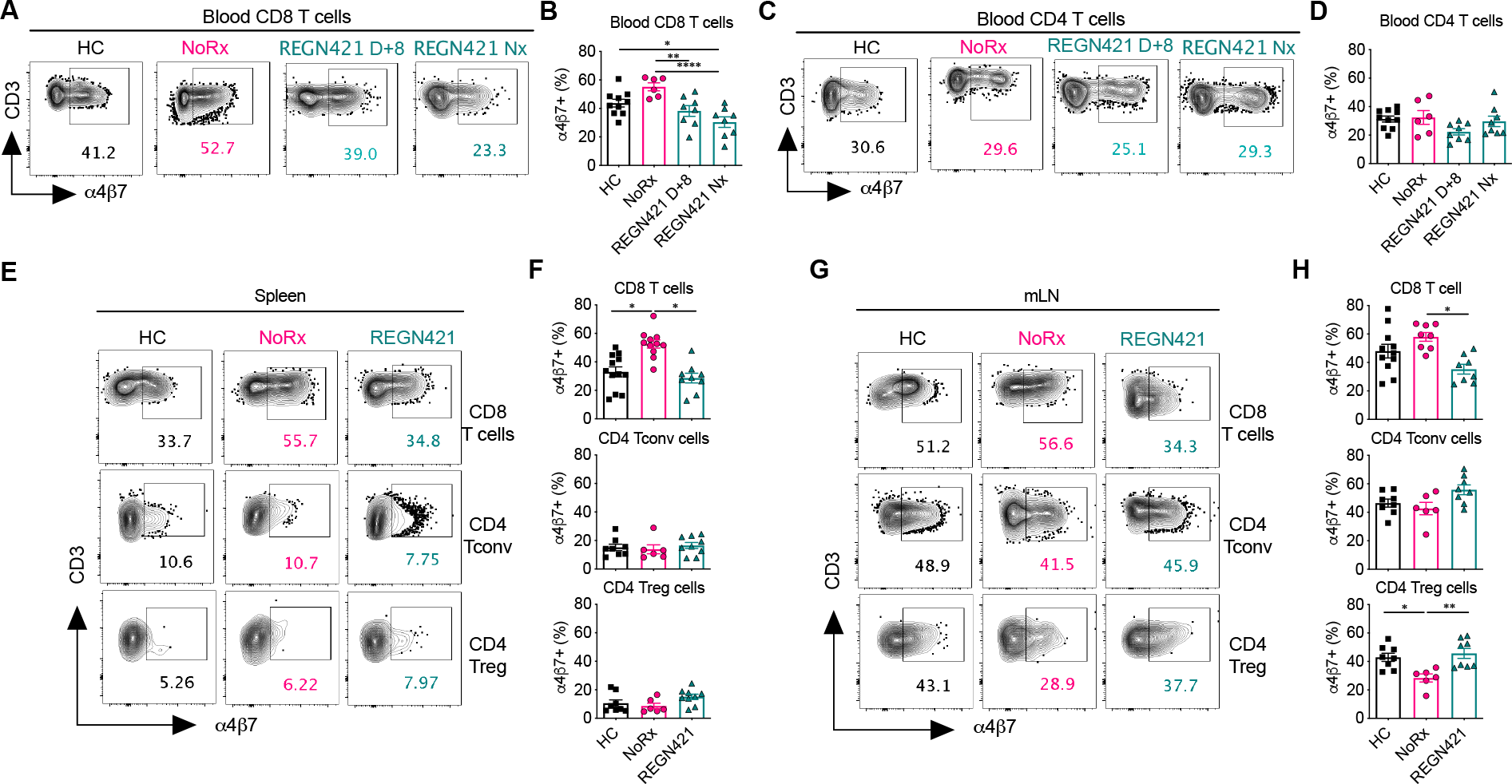
DLL4 blockade specifically prevents T cell expression of the gut-homing a4b7 integrin during GVHD. (A-D) Representative flow cytometry plots (A,C) and data (B,D) depicting the relative number of a4b7^+^CD8^+^ and a4b7^+^CD4^+^ T cells in the blood of healthy controls (n=10) and allo-HCT recipients from the NoRx aGVHD (n=6) and REGN421 (n=10) cohorts. Data from the REGN421 cohort was collected at terminal analysis and day +8 (time-matched with the NoRx cohort). *p<0.05, **p<0.01, ***p<0.001, one-way ANOVA with Tukey post-hoc-test. (E-H) Representative flow cytometry plots (E,G) and data (F,H) depicting the relative number of a4b7^+^CD8^+^, a4b7^+^ conventional CD4^+^ and a4b7^+^ regulatory CD4^+^ T cells in spleen and mesenteric lymph nodes of healthy controls (n=8-12 depending on the organ) vs. allo-HCT recipients from the NoRx aGVHD (n=8-11 depending on the organ) or REGN421 (n=10) experimental cohorts. *p<0.05, **p<0.01, one-way ANOVA with Tukey post-hoc-test.

To test if regulation of a4b7 cell surface levels in REGN421-prophylaxed animals correlated with Tcon and Treg abundance in the GI tract compared to NoRx controls, we studied gut samples from transplant recipients (Fig. 4). In the NoRx aGVHD cohort, we observed increased infiltration by activated Ki67^+^CD3^+^ T cells in the intestine at terminal analysis (Fig. 4A-C, Fig. S3F) compared to healthy controls. DLL4 blockade decreased colonic infiltration by Ki67^+^ T cells (Fig. 4A-C, Fig. S3F). In addition, REGN421 increased the Treg:Tcon ratio and the percentage of CD25^+^CD127^−^FoxP3^+^ Treg in the GI tract compared to the NoRx aGVHD cohort (Fig. 4D-E). In the liver, however, the Treg:Tcon ratio was similar in REGN421-treated and NoRx cohorts (Fig. 4D-E). The differential impact of REGN421 on Treg and Tcon recruitment to the GI tract could be related to the relatively preserved abundance of a4b7 in Treg after DLL4 blockade as compared to the reduced cell surface levels of a4b7 in Tcon (Fig. 3E-H).

**Figure 4.**
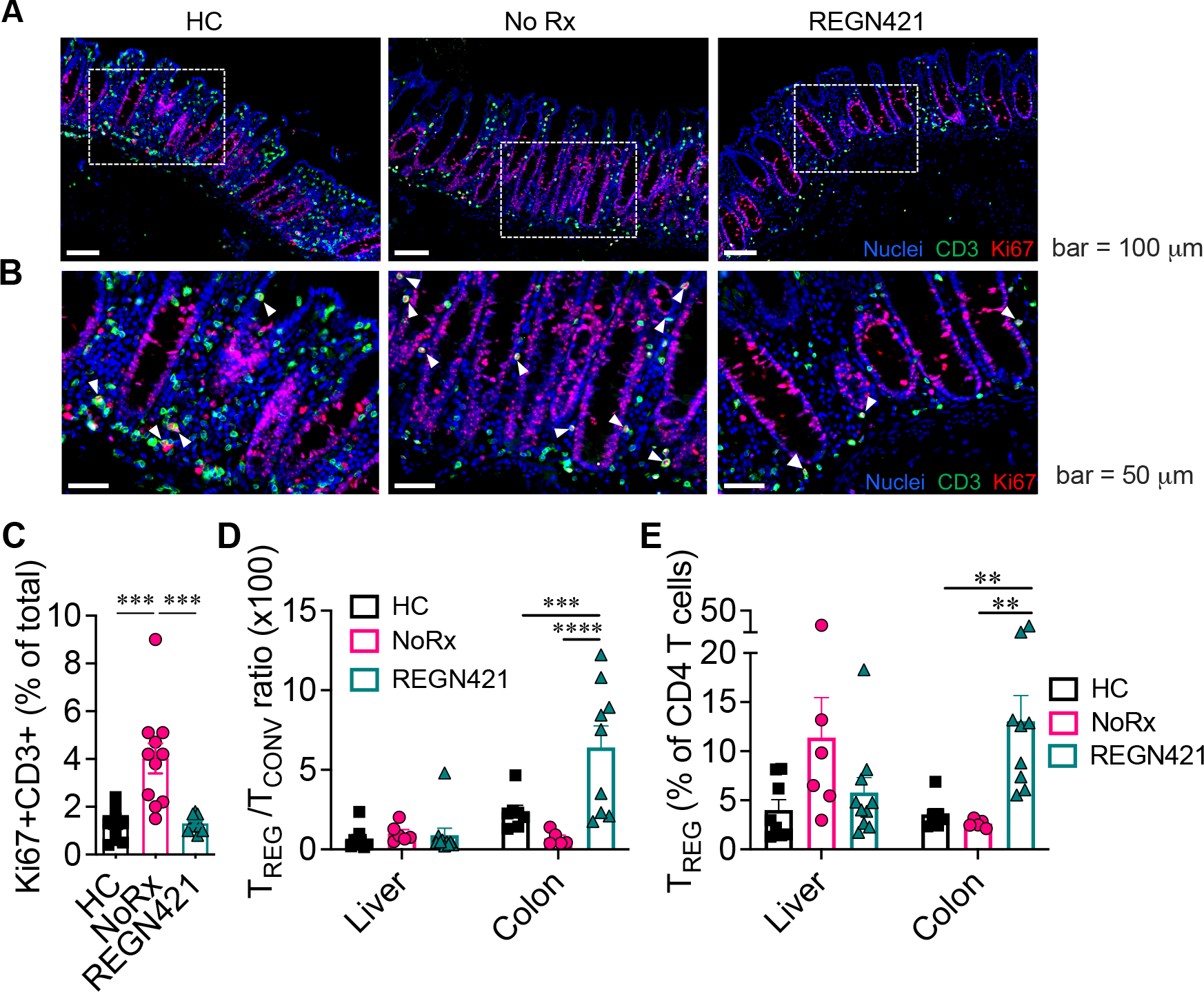
DLL4 blockade inhibits the accumulation of activated tissue-infiltrating T cells in the intestine during aGVHD. (A-C) Immunofluorescence microcopy of paraffin-embedded colon collected at terminal analysis in different experimental cohorts. Samples were from healthy controls (HC) vs. terminal analysis of allo-HCT recipients from unprophylaxed aGVHD (NoRx) or REGN421-treated experimental cohorts (aDLL4). (A) Representative staining for CD3 (green), Ki67 (red) and nuclei visualized by Hoechst (blue). (B) Cropped areas in (A) are enlarged with white arrowheads pointing to CD3^+^Ki67^+^ T cells. (C) Quantification of Ki67^+^CD3^+^ T cells among total nucleated cells (n=11 images from 5 animals per group). ***p<0.001, Kruskal-Wallis multiple comparison test. (D) Treg:Tcon ratio and (E) % of Foxp3^+^Treg among CD4^+^ T cells in the liver and colon from healthy controls (n=8) versus at terminal analysis in allo-HCT recipients from unprophylaxed aGVHD (NoRx, n=7) or REGN421-treated (n=10) cohorts. ***p<0.001, ****p<0.0001, one-way ANOVA with Tukey post-hoc-test.

### In a mouse allo-HCT model, Notch inhibition blunts a4b7 upregulation and intestinal accumulation of conventional CD4^+^ and CD8^+^ T cells, but preserves intestinal Treg

To gain mechanistic information about T cell-intrinsic effects of Notch on cell surface a4b7 and gut homing, we used a lethally irradiated parent-to-F1 MHC-mismatched mouse model with a sublethal dose of T cells, blocking all Notch signals in donor CD4^+^ and CD8^+^ T cells through expression of the pan-Notch inhibitor DNMAML (Fig. 5A). These experiments enabled a detailed kinetic analysis of a4b7 expression and T cell homing to the gut that was not possible in NHP. DNMAML expression in T cells prevented both weight loss and thymic injury, two representative features of aGVHD (Fig. S4A-B). Moreover, and as reported previously (*19,21, 24*), Notch inhibition led to an increased proportion of Tregs at multiple sites in HCT recipients (Fig. S4C-D). Next, we quantified cell surface a4b7 in activated donor-derived T cell subsets at day 4, 7, 14 and 28 after transplant. At day 4, DNMAML blunted the induction of cell surface a4b7 in CD8^+^ and in Foxp3^−^CD4^+^ Tcon in the spleen and mLN (Fig. 5B-C). In contrast, Foxp3^+^CD4^+^ DNMAML Treg had preserved or even increased a4b7 on their surface. The impact of Notch inhibition on a4b7 levels persisted at later time points in CD8^+^, but not CD4^+^ T cells, similarly to our findings in NHPs and showing evolutionary conservation (Fig. 5B-C, Fig. 3).

**Figure 5.**
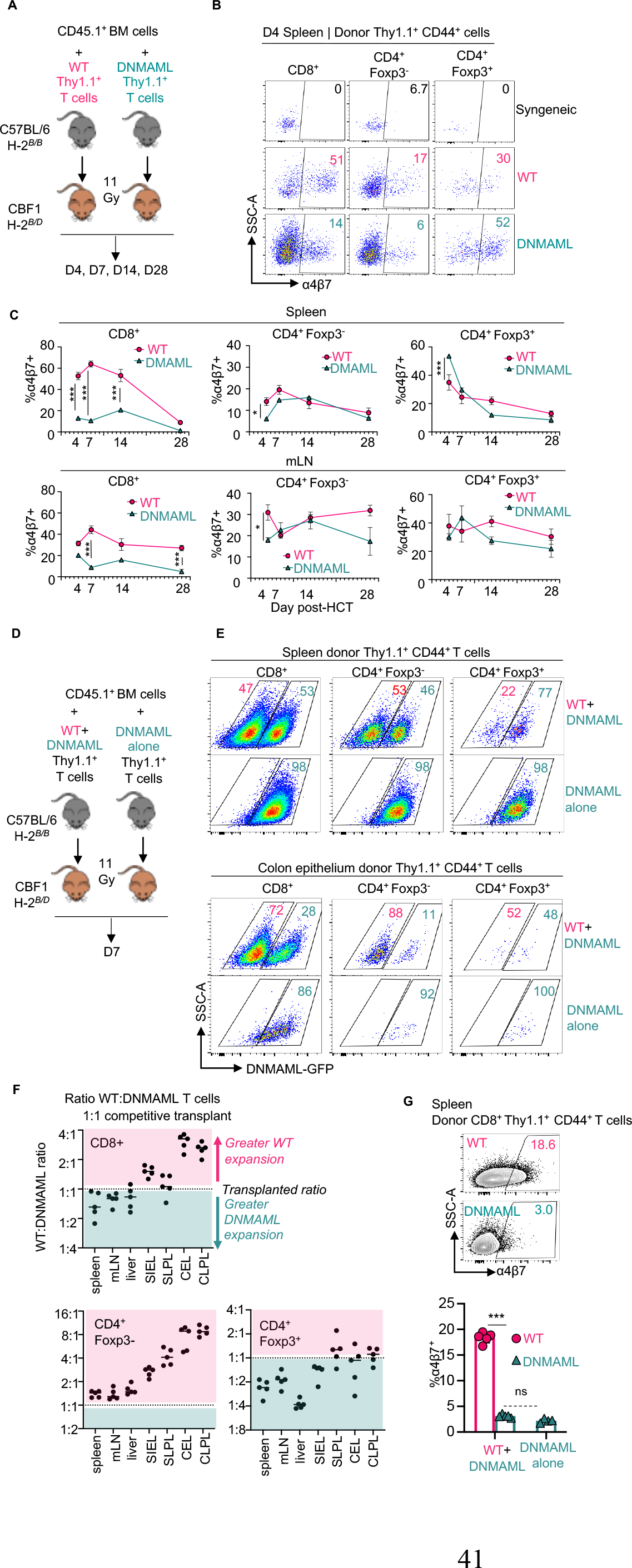
Cell-intrinsic canonical Notch signals control cell surface α4β7 and gut-homing potential in alloreactive T cells. (A) 1x10^6^ Thy1.1^+^ syngeneic CBF1 T cells (H-2^b/d^), allogeneic wild-type B6 T cells (H-2^b/b^) vs. allogeneic Notch-deprived B6-DNMAML T cells (H-2^b/b^) were transplanted into lethally irradiated CBF1 recipients (H-2^b/d^). Recipients received CD45.1^+^Thy1.2^+^ T cell-depleted BM to distinguish BM-derived cells from the T cell inoculum. Representative flow cytometric analysis of α4β7 in donor-derived CD44^+^CD8^+^, Foxp3^−^CD4^+^ and Foxp3^+^CD4^+^ T cells in the spleen at day 4 (B) and summary data at indicated times post- transplant in spleen and mLN (C). n=3 mice per group and time point. (D) CBF1 recipients were transplanted as in (A), but with 1:1 mixed wild type:DNMAML donor T cells vs. DNMAML T cells alone. (E) Representative flow cytometric plots showing relative accumulation of wild type compared to DNMAML-GFP^+^ T cells in spleen (top) vs. colon epithelium (bottom) at day 7. (F) Organ-specific accumulation of wild type to DNMAML T cells from the 1:1 competitive transplant was calculated for each T cell subset and expressed as greater than (red) or less than (green) the initial ratio. (G) Representative flow cytometric analysis of α4β7 in donor-derived CD8^+^ T cells from inocula of wild type and DNMAML T cells (1:1, left) vs. DNMAML T cells alone. *p<0.05, ***p<0.001, one-way ANOVA with Tukey’s post-hoc-test. Syn = syngeneic. mLN = mesenteric lymph node, SI = small intestine, C = colon, EL = epithelial lymphocytes, LPL = lamina propria lymphocytes.

Because Notch inhibition had divergent impact on a4b7 abundance in Tcon versus Treg that could differentially affect homing to the GI tract, we designed a competitive assay for the relative accumulation of wild-type and Notch-deprived DNMAML T cell subsets in recipient organs (Fig. 5D). We injected a 1:1 mixture of wild type and DNMAML T cells into irradiated allo-HCT versus DNMAML T cells alone as control. We tracked the percentage of Notch- deprived T cells in secondary lymphoid organs and GVHD target organs based on expression of the DNMAML-GFP fusion protein in donor-derived CD44^+^ T cells (Fig. 5E-F). In the spleen, mLN and liver, we recovered a ratio of wild type to DNMAML Tcon close to the 1:1 ratio in the inoculum, both for CD8^+^ and Foxp3^−^CD4^+^ Tcon. In contrast, the colon (Fig. 5E-F) and small intestine (Fig. 5F) contained a markedly decreased proportion of DNMAML-GFP^+^CD8^+^ and Foxp3^−^CD4^+^ T cells, consistent with cell-autonomous effects of Notch inhibition on their accumulation in the gut. For Foxp3^+^CD4^+^ Treg, however, we observed expansion of DNMAML- GFP^+^ Treg in secondary lymphoid organs, and preservation of DNMAML Treg accumulation in the gut (as evidenced by a lower wild type to DNMAML ratio among Treg compared to Tcon) (Fig. 5E-F). Similar findings were apparent with a 1:2 wild-type to DNMAML T cell inoculum (Fig. S4E). Using the same design, we confirmed that DNMAML-mediated Notch inhibition decreased cell surface a4b7 via cell-autonomous mechanisms (Fig. 5G, Fig. S4F-G). Thus, Notch signals increase α4β7 abundance and gut-homing potential in alloreactive Tcon, but not in Treg. Altogether, complementary NHP and mouse data suggest that Notch inhibition blunts intestinal homing of Tcon, while preserving that of Treg, leading to an increased Treg/Tcon ratio in the GI tract early after transplantation.

### DLL4 blockade induces complex changes in the T cell transcriptome and is associated with normalization of an aGVHD-specific T cell GI-infiltration signature

To gain insights into how REGN421 affects T cells after allo-HCT, we explored transcriptomic changes using Rhesus macaque-specific gene arrays (*12, 13, 15, 16*) (Table S1). T cells were purified from blood on day 15 after HCT (‘REGN421.D15’ group) and at time of necropsy (‘REGN421.Nx’ group). Transcriptomes were compared to three other datasets: (1) the NoRx aGVHD cohort, with gene arrays obtained at necropsy (MST = 8 days, n=12); (2) recipients prophylaxed with the KY1005 anti-OX40L antibody (MST = 19.5 days, blood for gene arrays obtained at day 15, n=4) (*13*); (3) recipients prophylaxed with the FR104 anti-CD28 (MST = 26 days, blood for gene arrays obtained at day 15, n=3) (*12*). Importantly, neither OX40L nor CD28 blockade offered specific protection from clinical GI-aGVHD, and pathological scores at terminal analysis were not different in KY1005, FR104 and NoRx aGVHD cohorts (Fig. S2D).

We first sought to identify a blood T cell transcriptomic signature associated with inhibition of GI infiltration after DLL4 blockade. Compared to NoRx aGVHD controls, blood T cells from REGN421-treated animals at day 15 and at necropsy demonstrated decreased enrichment for an aGVHD-specific GI infiltration signature recently discovered by our group (*11*), which includes multiple chemotaxis-related genes. In contrast, this signature was enriched in the NoRx vs. healthy control cohort (Fig. 6A). Notably, neither anti-OX40L nor anti-CD28- prophylaxed cohorts normalized the T cell GI infiltration signature (Fig. 6A), suggesting that these interventions had effects distinct from those of DLL4 blockade.

**Figure 6.**
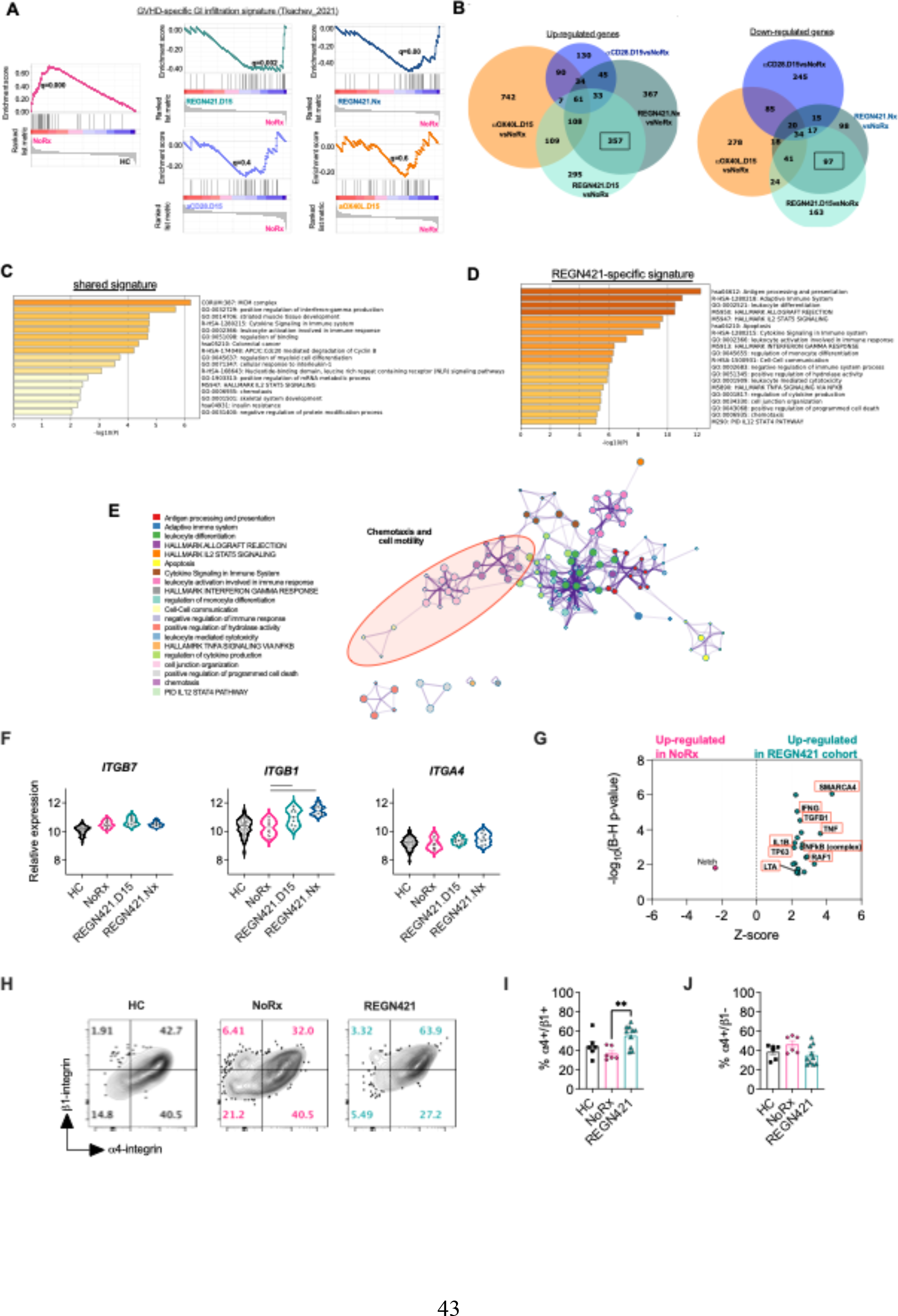
DLL4 blockade induces unique changes in the T cell transcriptome and is associated with normalization of a GVHD-specific T cell GI-infiltration signature. (A) GSEA plots showing enrichment for a recently defined GVHD-specific GI infiltration signature (*11*) in unprophylaxed aGVHD (NoRx) compared to healthy controls (HC) and depicting normalization of this signature upon DLL4 inhibition, but not OX40L or CD28 blockade. (B) Venn diagram depicting the overlap of differentially expressed (DE) genes for each aGVHD monoprophylaxis cohort in comparison with NoRx aGVHD. (C-D) Functional annotation and pathway enrichment analysis was performed with 95 DE genes (61 over-represented, 34 under- represented) that were affected by the three GVHD monoprophylaxis protocols in comparison with the NoRx aGVHD (the “shared signature”) (C) and with 454 DE genes (357 over- represented, 97 under-represented) that were affected only by DLL4 blockade (the “REGN421- specific signature”) using the Metascape tool (Metascape.org). Top enriched ontology clusters are shown for each signature. (E) Visualization of top enriched functional pathways, as analyzed by hierarchical clustering and visualized using Metascape. (F) Relative abundance of *ITGA4*, *ITGB7* and *ITGB1* mRNA in T cells from healthy controls (HC), unprophylaxed allo-HCT recipients (NoRx) and allo-HCT recipients receiving anti-DLL4 antibodies (REGN421.D15: blood T cells at day 15; REGN421.Nx: blood T cells at necropsy). Lines indicate statistically significant differences between cohorts (p<0.05 using a Benjamini-Hochberg correction). (G) The 454 DE genes specifically affected by anti-DLL4 were analyzed using Ingenuity Pathway Analysis to identify upstream regulators activated or inhibited in REGN421 vs. NoRx aGVHD cohorts. Shown are positive upstream regulators of *ITGB1* expression. (H-J) Representative flow cytometry plots (H) and data (I-J) depicting the proportion of spleen CD8^+^ T cells with cell surface a4 and b1 integrins, versus a4 in the absence of b1 in healthy control (HC, n=6), NoRx aGVHD (n=6) and REGN421 (n=10) cohorts. **p<0.01, one-way ANOVA with Tukey post-hoc-test

To better understand REGN421-specific mechanisms of aGVHD protection, we identified differentially expressed genes that were exclusive to the comparison of REGN421 versus NoRx, and not found when comparing anti-OX40L or anti-CD28 to NoRx (Fig. 6B, Table S2-S3). We identified 357 overrepresented and 97 underrepresented transcripts specific to the REGN421 versus NoRx comparison. In contrast, we identified only 61 overrepresented and 34 underrepresented shared transcripts among all three interventions compared to NoRx (Fig. 6B, Table S4). Functional annotation and pathway analysis performed using Metascape (*41*) revealed that the shared GVHD protection signature predominantly involved cell cycle/DNA replication as well as cytokine signaling pathways (Fig. 6C, Table S5). Notably, the REGN421-specific GVHD-protection signature was enriched in cell-cell communication and chemotaxis-related genes (Fig. 6D-E, Table S6).

### DLL4 blockade increases expression of the integrin b1 subunit while decreasing integrin b7 on the T cell surface

We next investigated how Notch inhibition affects cell surface abundance of the a4b7 gut-homing integrin after allo-HCT in NHP T cells. *ITGA4* and *ITGB7* transcripts (encoding subunits of the a4b7 integrin) were *not* decreased by DLL4 blockade with REGN421 (Fig. 6F). Thus, a simple transcriptional effect did not account for decreased a4b7 in T cells from REGN421-treated recipients. In contrast, *ITGB1* mRNA was increased in T cells from the REGN421 cohort (Fig. 6F). *ITGB1* encodes integrin b1, which interacts with the a4 integrin chain and has been reported to compete with b7 for a4 binding (*42*). To further investigate upregulated *ITGB1* expression in the REGN421 versus NoRx cohorts, ingenuity pathway analysis was utilized to identify putative upstream regulators. We found 22 upstream signaling molecules/pathways activated by REGN421, of which 9 included *ITGB1* as a target gene (Fig. 6G, Table S7).

We next evaluated surface abundance of b1, b7 and a4 integrin chains. Consistent with mRNA, overall cell surface levels of b1 integrin were increased in CD8^+^ T cells from REGN421-treated as compared to NoRx animals (Fig. S5). In contrast, surface b7 integrin was decreased (Fig. S5), despite comparable *ITGB7* transcripts in both cohorts (Fig. 6F). When assessing cell surface a4 and b1 together, we observed an increased proportion of a4^+^b1^+^ CD8^+^ T cells in the REGN421-treated as compared to NoRx recipients (Fig. 6H-I). Moreover, nearly all b1^+^CD8^+^ T cell simultaneously stained for a4, and the proportion of a4^+^b1^−^ T cells was decreased (Fig. 6H-J). These data are consistent with inhibition of b7 integrin trafficking to the cell surface via competitive binding of a4 by b1 (*42, 43*). Cell surface a4b7 is also regulated by availability of a4 integrin (*44*). However, *ITGA4* mRNA was similar in T cells from REGN421- prophylaxed and NoRx cohorts (Fig. 6F) and the proportion of a4^+^CD8^+^ T cells was actually increased in the REGN421 cohort (Fig. S5). Thus, we hypothesize that DLL4 blockade results in upregulated expression of the integrin b1 subunit, leading to increased competition for a4 binding and decreased abundance of the a4b7 heterodimer.

### Loss of integrin b1 rescues α4β7 expression in Notch-deprived alloreactive T cells

To test if b1/b7 integrin subunit competition was responsible for the effects of Notch inhibition on cell surface a4b7 in T cells, we used a mouse allo-HCT model with genetic inactivation of *Itgb1* and Notch signaling in T cells (with *Cd4-Cre;Itgb1^f/f^;ROSA^DNMAML^* mice and respective controls as allogeneic donors). We lethally irradiated CBF1 recipients (H-2^b/d^) and transplanted them with allogeneic T cell-depleted bone marrow plus T cells from *Cd4-Cre;ROSA^DNMAML^*, *Cd4-Cre;Itgb1^f/f^*, *Cd4-Cre;ROSA^DNMAML^;Itgb1^f/f^* or wild-type littermate controls (on a mixed B6/129 background, all H-2^b^). In activated donor-derived CD44^hi^CD8^+^ T cells, pan-Notch inhibition with DNMAML increased the proportion of T cells co-expressing a4 and b1, and decreased the proportion of a4^+^b1^−^ T cells (Fig. 7A). Donor-derived CD4^+^ T cells showed similar findings (Fig. 7B, Fig. S6A-B). Importantly, the proportion of cells co-expressing cell surface a4 and b7 was lower among a4^+^b1^+^ than a4^+^b1^−^ T cells, consistent with b1/b7 competition for a4 binding (Fig. 7C). In activated donor CD44^hi^CD8^+^ T cells, pan-Notch inhibition with DNMAML decreased co-expression of a4 and b7 integrin subunits on the cell surface, and this effect was fully rescued by *Itgb1* inactivation (Fig. 7D). The same was true in donor-derived conventional Foxp3^−^CD4^+^ T cells (Fig. 7E, Fig. S6C-D). In contrast, Notch inhibition did not decrease cell surface a4b7 in Treg, and even increased it in the absence of *Itgb1* (Fig. 7E, Fig. S6C-D). To further corroborate these findings, we used an antibody specific for the a4b7 heterodimer (Fig. S6E). DNMAML decreased a4b7/LPAM-1 reactivity in CD8^+^ T cells, and this effect was rescued by *Itgb1* inactivation.

**Figure 7.**
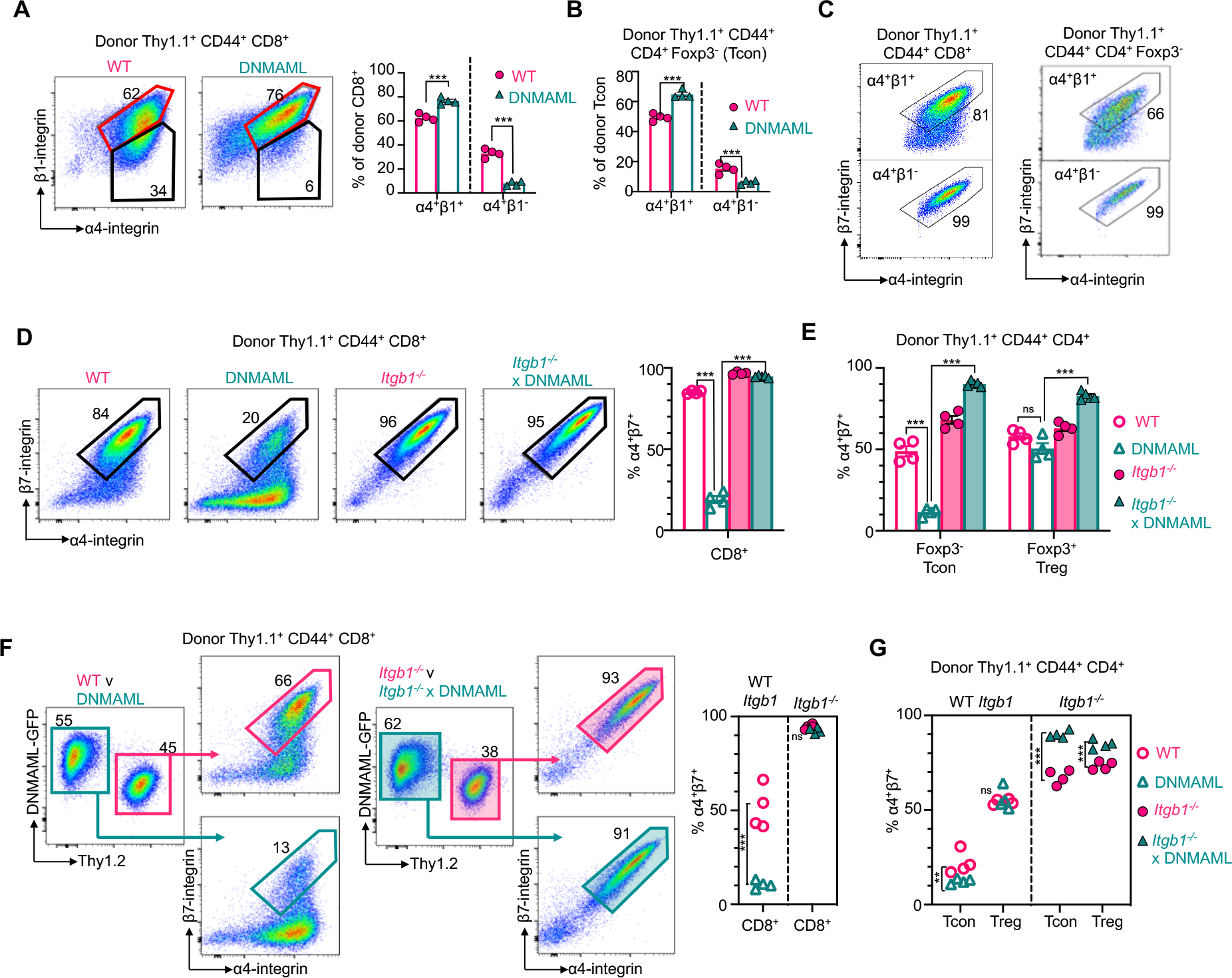
Loss of integrin b1 rescues α4β7 in Notch-deprived alloreactive T cells. (A-E) 20x10^6^ Thy1.1^+^ or Thy1.1/1.2^+^ B6 splenocytes and lymph node cells from mice with wild type (WT), Notch-deprived (DNMAML), integrin b1-deficient (*Itgb1^-/-^)* or Notch-deprived and integrin b1-deficient (*Itgb1^-/-^* x DNMAML) T cells were transplanted into lethally irradiated CBF1 recipients (Thy1.2^+^, H-2^b/d^) along with 1x10^6^ T-cell depleted Thy1.2^+^ BM cells. Spleens and mLNs were harvested on day 4.5. (A) Representative flow cytometry plots and summary data for α4β1 in donor-derived CD44^+^CD8^+^ T cells (spleen, day 4.5). (B) Summary data for α4β1 in donor-derived CD44^+^Foxp3^−^CD4^+^ conventional T cells (Tcon). (C) Flow cytometry plots for α4β7 in α4^+^β1^+^ (top) and α4^+^β1^-^ (bottom) wild type CD8^+^ (left) and CD4^+^ Tcon (right). (D) Flow cytometry plots and summary data for α4β7 in donor CD8^+^ T cells, and summary data (E) in CD4^+^Tcon and Foxp3^+^CD4^+^ regulatory T cells (Treg). (F-G) CBF1 mice were transplanted as in (A-E), but with mixed 15x10^6^:15x10^6^ wild type:DNMAML T cells, or 15x10^6^:15x10^6^ *Itgb1^-/-^*:*Itgb1^-/-^* DNMAML T cells. (F) Flow cytometry plots showing α4β7 in wild type vs. DNMAML CD8^+^ cells (left) and *Itgb1*^-^*^/-^* vs. *Itgb1^-/-^* x DNMAML CD8^+^ T cells (right) as well as summary data (far right). (G) Cell surface α4β7 in competing CD4^+^Tcon and CD4^+^Tregs. n=4 mice per group. **p<0.01, ***p<0.001, two-way ANOVA with Tukey’s post- hoc-tests. ns = not significant.

Finally, we asked whether the *Itgb1*-dependent effects of Notch inhibition in T cells were cell-autonomous. To this end, we co-transplanted wild-type and DNMAML T cells with or without *Itgb1* expression into allogeneic recipients (Fig. 7F-G, Figure S6F-G). As assessed via detection of individual a4 and b7 subunits as well as LPAM-1 reactivity, DNMAML markedly decreased the proportion of T cells with cell surface a4b7 integrin heterodimers through cell-autonomous mechanisms, and this effect remained dependent on *Itgb1* expression in Tcon. Thus, Notch inhibition blunted the abundance of cell surface a4b7 in an *Itgb1*-dependent manner in CD4^+^ and CD8^+^ Tcon after allo-HCT. In contrast, expression of this critical gut-homing integrin on the cell surface was preserved in Notch-deprived Treg (Fig. S7).

## DISCUSSION

Our observations uncover a critical impact of Notch signaling in the pathogenesis of GI-aGVHD, a major life-threatening complication of allo-HCT. Complementary investigations in mouse and NHP models demonstrated dominant effects of the Notch ligand DLL4 in GVHD that were highly conserved during evolution and could be targeted therapeutically. Furthermore, DLL4/Notch inhibition protected mice and NHP from GI-aGVHD through unique mechanisms that differed from those of other, more global inhibitors of T cell activation (*12, 13*). This mechanism included the normalization of a transcriptional T cell gut-homing signature recently discovered in the NHP aGVHD model (*11*). Decreased cell surface abundance of the a4b7 gut- homing integrin was a prominent and conserved effect of DLL4/Notch inhibition in effector T cells, whereas its expression was maintained in Tregs (Fig. S7). Altogether, these findings highlight new therapeutic opportunities and provide new insights into evolutionary conserved mechanisms of GI infiltration and GVHD pathogenesis that are predicted to also operate in human patients.

First discovered in Drosophila and studied extensively for its multiple functions in metazoans, Notch signaling operates as a ligand/receptor system via biochemical mechanisms that have been conserved during evolution (*23*). Yet, mammals harbor multiple Notch ligands and receptors that are deployed in specific contexts, and whose relative importance can drift between species (*22, 45, 46*). Thus, observations in mice should not be assumed to apply strictly to humans, even if the Notch pathway as a whole exerts conserved biological functions. This limitation is important for translational investigations, as it is not typically addressed by preclinical work in small animal models. In contrast, our findings in mice and NHP document conserved effects of the Notch ligand DLL4 on T cell infiltration into the GI tract and GI aGVHD pathogenesis, with high single agent activity of DLL4 inhibitors. DLL4 is one of four agonistic Notch ligands in mammals, together with DLL1 and JAGGED1/2 family ligands. We previously reported a critical role for Delta-like ligands in multiple mouse models of GVHD, with a dominant role for DLL4 and a more minor role for DLL1 (*18, 19, 21, 24, 25*). Our data now indicate that the unique immunological functions of DLL4 in GVHD have been conserved during evolution. The relative additional importance of DLL1 will require further investigation.

Furthermore, our findings support a major role for Notch ligands of the Delta-like but not Jagged family in immune regulation.

Beyond the conserved role of DLL4 among other Notch ligands, key immunological effects of the Notch pathway in aGVHD were conserved during evolution. Both in mice and in NHP, DLL4/Notch inhibition did not protect from GVHD through global immunosuppression. Analysis of T cell reconstitution in REGN421-treated recipients showed expansion of donor-derived T cells with central memory and effector memory phenotypes, while naïve T cells were depleted, suggesting preserved alloantigen-mediated T cell activation and differentiation (*47, 48*). In contrast, anti-CD28 and anti-OX40L blunted in vivo T cell activation after allo-HCT in NHP (*12, 13*), an effect that has also been observed in clinical transplantation (*31*) and may lead to more global immune suppression. In mice, we reported preserved in vivo expansion and differentiation of Notch-deprived alloreactive T cells in secondary lymphoid organs despite profound protection from aGVHD (*18, 19, 21*). Importantly, Notch inhibition preserved efficient graft-versus-tumor (GVT) effects (*19, 21*). Although GVT activity cannot be studied directly with existing NHP models, our findings predict that DLL4/Notch blockade should preserve beneficial systemic anti-infectious and anti-cancer T cell immune responses after allo-HCT, which is highly desirable in patients. In addition, a short duration of DLL4/Notch blockade was sufficient to provide protection from aGVHD, with all benefits conferred by a single dose of antibodies at time of transplantation. Thus, any subsequent effects of the Notch pathway in T cell immunity should be restored after clearance of the blocking antibodies.

Instead of global immunosuppression, DLL4/Notch inhibition provided strikingly specific protection from GI-aGVHD, an effect not observed with single agents tested so far for aGVHD prevention in NHP. The NHP model coupled to mechanistic investigations in mice provided unique, fundamental insights into conserved mechanisms of T cell infiltration into the GI tract after allo-HCT. We speculate that rapid exposure of donor T cells to Notch ligands in secondary lymphoid organs within days after allo-HCT induces a gut-homing program that promotes early seeding of the gut by alloreactive T cells. The a4b7 integrin dimer was previously reported to play an important role in T cell migration/infiltration into the gut and in the pathogenesis of intestinal GVHD in mice (*36, 49*). a4b7 interacts with its ligand, MAdCAM1, expressed by endothelial cells in the intestinal lamina propria and in gut-associated lymphoid tissues (*50, 51*). Hanash and collaborators reported a critical role for this interaction in T cell recruitment to a vulnerable compartment containing intestinal stem cells at the base of intestinal crypts, with early injury to these cells playing a critical role in propagating GVHD (*5*). Both in mice and in NHP, DLL4/Notch inhibition decreased cell surface a4b7 in conventional effector T cells. Mechanistically, these effects were not related to transcriptional regulation of *Itga4* or *Itgb7* gene expression by Notch, but instead to upregulated expression of *Itgb1* encoding the integrin subunit b1. Through this unique mechanism, b1 appeared to compete with b7 for a4 binding as reported in other contexts (*42*). Indeed, the presence of *Itgb1* was necessary for Notch inhibition to decrease cell surface a4b7 in T cells after mouse allo-HCT. Importantly, upregulated *Itgb1* expression was detected in Notch-deprived NHP T cells together with normalization of a unique T cell gut-homing signature previously identified in NHP through unbiased transcriptomic analysis (*11*). Thus, Notch emerges as a central regulator of T cell recruitment to the gut after allo-HCT, and Notch inhibitors exert beneficial effects in GVHD through effects that other interventions do not recapitulate.

As a unique feature of DLL4/Notch inhibition in mice and NHP, cell surface a4b7 decreased in CD4^+^ and CD8^+^ Tcon early after allo-HCT while being maintained or even slightly increased in Treg. In addition, Notch inhibition decreased the accumulation of Tcon in the gut while preserving Treg recruitment, resulting in an increased Treg:Tcon ratio. How Notch signaling differentially regulates a4b7 expression in Tcon and Treg remains to be discovered. However, it may be essential in shifting the early balance of T cells in the gut, creating a higher Treg to Tcon ratio and a microenvironment with local immunosuppressive properties that blocks initiation of intestinal GVHD. Therapeutically, pan-inhibition of a4b7 is being explored in GVHD, e.g. with the antibody vedolizumab (*52, 53*). A limitation of this intervention is to inhibit a4b7 equally in Tcon and in Treg. In contrast, the differential effects of DLL4/Notch inhibition in Tcon and Treg may preserve Treg recruitment, tilting the local ratio of pathogenic and protective T cells at a crucial stage in which the GI tract is seeded by T cells after allo-HCT. In addition, DLL4/Notch inhibition may exert protective effects independent of a4b7 regulation as part of its broader impact on the T cell gut homing signature that we have identified.

For the first time, our work provides important information about key safety indicators of DLL4 inhibitors in a model closely recapitulating human allo-HCT. This is essential, given that the complex peri-transplant environment is not replicated in mice. Understanding the safety profile of REGN421 prophylaxis is even more important since, in allo-HCT, administration of Notch inhibitors coincides with a period of recovery from pre-transplant conditioning, a time of heightened clinical risk for patients. Reassuringly, short-term systemic DLL4 blockade with REGN421 did not trigger unexpected side effects in our NHP model, while preserving rapid engraftment as well hematopoietic and immune reconstitution. This is significant, since unlike DLL4 blockade, pan-Notch inhibition with g-secretase inhibitors or other non-selective approaches were poorly tolerated after allo-HCT in mice due to Notch inhibition in intestinal crypts (*21*). This intestinal toxicity is avoided by selective DLL4 blockade alone, which is not sufficient to trigger defects in intestinal homeostasis (*21, 54*). Furthermore, a single dose of REGN421 was sufficient to confer protection from GI-aGVHD, thus avoiding concerns about long-term DLL4 blockade which was reported to cause cardiovascular side effects after months of cumulative inhibition (*29*).

Altogether, our data provide new fundamental and translational insights into the role of Notch signaling in T cell pathogenesis during aGVHD, with complementary input from observations in mice and NHP. We reveal for the first time a central role for Notch signaling in the selective regulation of T cell homing and retention in the gut, with a differential impact on conventional T cells as opposed to Tregs. Practically, the unique single agent activity of REGN421 in NHP nominates DLL4 blockade as a worthwhile strategy to develop for the prevention of GI-aGVHD in patients, especially since it is predicted to have activity on a pathogenic gut homing program that other interventions do not neutralize. Possible future strategies include the combination of DLL4 inhibitors with other agents to blunt aGVHD at non-intestinal sites. Alternatively, it would be attractive to deploy DLL4 blockade in the setting of transplantation protocols in which GI involvement is the dominant manifestation of aGVHD. In addition, it will be interesting to investigate if DLL4/Notch signaling exerts conserved effects on immune cell homing and retention to target organs in other immunological contexts, such as intestinal autoimmune disorders.

## MATERIALS AND METHODS

### Non-human primates

Healthy, immunologically naïve juvenile and adult (mean age 5.9 years, range 2.1-14.8) Indian Rhesus macaques (Macaca Mulatta) of both sexes were obtained from breeding colonies at Alphageneis Inc., UC Davis National Primate Research Center or Washington National Primate Research Center (WNPRC), and were housed at WNPRC or at the Biomere Biomedical Research Models (BBRM). The study was conducted in strict accordance with USDA regulations and the recommendations in the Guide for the Care and Use of Laboratory Animals of the National Institutes of Health and was approved by the BBRM and the University of Washington/WNPRC Institutional Animal Care and Use Committees.

### NHP Study Design and Transplant Strategy

This was a prospective study in NHP designed to determine the biological role of Notch signaling in T cells during aGVHD, and the therapeutic effect of inhibiting Notch signaling via blocking the Notch ligand Delta-like ligand 4 (DLL4). Several cohorts of transplant recipients were studied: (1) Allogeneic transplants with no GVHD prophylaxis (abbreviated as ‘NoRx’, n = 11); (2) Allogeneic transplants receiving one dose of anti-DLL4 blocking antibodies REGN421 (Regeneron, Tarrytown, NY) (abbreviated as ‘REGN421’, n = 7); (3) Allogeneic transplants receiving three doses of REGN421 antibodies (n = 4). Animal demographic parameters, transplant characteristics, and doses of REGN421 antibodies administered in vivo are shown in Table S1.

This study used rhesus macaques that were housed at BBRM or at the Washington National Primate Research Center. Both REGN421x1 and REGN421x3 cohorts used half-sibling MHC haplo-identical donor and recipient pairs (Table S1). Experiments were performed using our previously described strategy for allogeneic HCT in rhesus macaques (*13–16*). Briefly, apheresis was performed after G-CSF mobilization (Amgen, 50 mcg/kg for 5 days), and an unmanipulated apheresis product was transplanted into allo-HCT recipients. The transplanted total nucleated cell dose (TNC) and CD3^+^ T cell dose are shown in Table S1. The pre-HCT preparative regimen consisted of total body irradiation (TBI) of 10.4 cGy given in two fractions per day for two days. Irradiation was delivered using either a linear accelerator Varian Clinac 23EX with a tissue-adjusted dose rate of 7 cGy/min or using gamma-irradiation with a Cobalt-60 isotope at a tissue-adjusted dose rate of 5.5 cGy/min. All transplants were performed with a central venous catheter placed for the length of the experiment. Antibacterial prophylaxis in all transplant recipients included Vancomycin (with a target serum concentration of 5-20 mcg/mL) and Ceftazidime, administered via central catheter, and Enrofloxacin (‘Baytril’), administered IM to all recipients with neutrophil counts less than 500 cells/μL. Antiviral prophylaxis (acyclovir, 10 mg/kg IV daily; cidofovir, 5 mg/kg IV weekly) and antifungal prophylaxis (fluconazole 5mg/kg oral or IV, given daily) were also employed. Leukoreduced (LRF10 leukoreduction filter, Pall Medical) and irradiated (2200 rad) platelet-rich plasma or whole blood was given for a peripheral blood platelet count of ≤50 × 10^3^ per μL or a hemoglobin <9 g/dL, respectively, or if clinically significant hemorrhage was noted. Blood product support adhered to ABO antigen matching principles.

The aGVHD clinical score was assessed daily and summarized weekly for allo-HCT recipients, as previously described (*13–16*). Briefly, the aGVHD clinical score increases with cumulative GI-specific abnormalities (diarrhea), liver-specific abnormalities (hyperbilirubinemia) and skin-specific abnormalities (extent and character of rash). It is important to note that the studies described here focused on the natural history of aGVHD that developed during prophylaxis, so that animals were not given supplementary treatment when GVHD was diagnosed. Rather, when pre-defined clinical endpoints were met (based on the BBRM and Washington National Primate Research Center veterinary standard operating procedures), animals were euthanized and a terminal analysis was performed. Thus, survival was directly related to the severity of clinical GVHD. Histopathologic scoring for GVHD was performed by an expert in GVHD histopathology (A.P.-M.) using a previously validated semi-quantitative scoring system (Grades 0.5-4). The pathologist was blinded to the treatment cohorts during the scoring process.

### GVHD prophylaxis regimens

Two DLL4 blockade strategies were evaluated. The first cohort of animals received one dose of GMP-grade anti-DLL4 blocking antibodies REGN421 (Regeneron) on day 0 at a dose of 3 mg/kg, 3 hours prior to infusion of the donor leukapheresis product. REGN421 (also called enoticumab) was developed and produced as a fully humanized IgG1 monoclonal antibody that binds human or NHP DLL4 (but not mouse DLL4) with sub-nanomolar affinity and inhibits DLL4-mediated Notch signaling, as described (*33*). The REGN421 dose was selected based on past pharmacokinetic measurements showing that it achieved therapeutic in vivo concentrations in healthy humans (*32*). The second cohort of animals received 3 mg/kg REGN421 on days 0, +7 and +14 relative to the HCT (the Day 0 dose was also given 3 hours prior to infusion of the leukapheresis product). The primary endpoint for REGN421-receiving cohorts was the proportion of animals surviving to 60 days post-transplant. No treatment for aGVHD was provided to recipients in this study; therefore, the clinical endpoints were unaffected by GVHD therapy. Pharmacokinetic studies of REGN421 were performed with blood obtained longitudinally from REGN421-treated recipients, using an ELISA developed at Regeneron (*32*).

### Chimerism determination

Whole blood, bone marrow and T cells (sorted by flow cytometry as CD3^+^CD20^−^) were analyzed for donor chimerism based on divergent microsatellite markers. Chimerism analysis was performed at the UC Davis veterinary genetics laboratory, as previously described (*14*).

### NHP necropsy and tissue processing

Animals were humanely euthanized followed by perfusion with sterile PBS (0.5 L/kg body weight), necropsy and tissue harvest. Blood PBMC were isolated using Ficoll-Paque PLUS gradient separation and standard procedures. Lymph nodes, spleen and bone marrow (from long bones) were mechanically disrupted by grinding through a metal strainer, followed by filtering through nylon 70 µm and 40 µm cell strainers. Lungs, liver, jejunum colon, and kidney samples were cut into small pieces, then digested using 150 KU DNase (Sigma-Aldrich) with 1 mg/mL Type I collagenase (Invitrogen) (lungs, liver and kidney) or 150 KU DNase (Sigma-Aldrich) with 50 mcg/mL Liberase TL (Roche; colon and jejunum) at 37°C for 1 hour with shaking, followed by passing through a metal strainer, and then 100 µm, 70 µm and 40 µm nylon cell strainers. The resulting cell suspension was further purified using a double-layer Percoll (GE Healthcare) gradient separation. The leukocyte fraction was collected at the interphase between

70% and 30% Percoll. Cells were either washed and immediately analyzed or were cryopreserved in 10% DMSO/90% FBS and stored in liquid nitrogen.

### Multiparameter flow cytometry on NHP samples

Samples were processed as previously described (*13*) and either analyzed fresh or from previously cryopreserved samples. Cryopreserved samples were washed with PBS and incubated with 200 µL 1:100 LIVE/DEAD Aqua (Invitrogen) for 20 minutes at 37°C using 96-well cell culture plates. After viability staining, cells were washed with staining buffer (PBS supplemented with 2% heat-inactivated FBS) and then stained with antibody cocktails against extracellular targets (Table S8) at 4°C for 20 minutes, followed by washing with the staining buffer. Fresh samples were washed with the staining buffer and directly stained with extracellular antibody cocktails without prior viability staining.

Samples stained with extracellular antibodies only were fixed with 1X BD Stabilizing Fixative and run within 24 hours after staining. Samples requiring intracellular staining were incubated in 250 µL Cytofix/Cytoperm solution (BD Biosciences) at 22°C for 20 minutes, followed by washing with 1X Perm/Wash buffer (BD Biosciences) and then incubated with intracellular antibody cocktails, mixed in 50 µL of the 1X Perm/Wash buffer, at 4°C for 30 minutes (Table S8). These samples were then washed and analyzed within 24 hours after staining. Samples requiring intranuclear staining were incubated in 250 µL Foxp3 fixation buffer (BioLegend) at 22°C for 20 minutes, followed by a washing step with Foxp3 wash buffer (BioLegend), and then incubated with intranuclear antibody cocktails, mixed in 50 µL of the Foxp3 wash buffer, at 4°C for 30 minutes (Table S8). These samples were then washed and analyzed within 24 hours after staining. Flow cytometry was performed on a BD FACS LSRFortessa and analyzed in FlowJo v.10.

### Flow Cytometric T cell sorting and microarray cohort designation

Using a FACSAria or FACSJazz Cell Sorter (BD), CD3^+^ T cells were sorted from the blood of 1) healthy controls, 2) allo-HCT recipients at day +15±1 post-transplant (as survival permitted), and 3) allo-HCT recipients at the time of terminal analysis. T cells were defined as CD3^+^CD20^−^ lymphocytes and were >90% pure based on post-sorting flow cytometric analysis, as previously described (*15, 16*).

### NHP microarray and data analysis

Following T cell purification, RNA was stabilized in T cell lysates with RLT buffer (Qiagen) supplemented with 1% (vol/vol) beta-mercaptoethanol (Sigma) and RNA was purified using RNEasy Column Kit (Qiagen). RNA was quantified using a Nanodrop Spectrophotometer (ThermoFisher Scientific) and purity was confirmed with an RNA 6000 Nano Kit (Agilent). The purified RNA was sent to the Vanderbilt Technologies for Advanced Genomics Core and to the Oregon Health Sciences University Gene Profiling Shared Resource, where RNA quantity and quality were verified, followed by cDNA/cRNA synthesis, and target hybridization to GeneChip Rhesus Macaque Genome Array (Affymetrix). The resultant fluorescent signals were processed and normalized using the Robust Multichip Averaging (RMA) Method (*55*). The microarrays were performed in 10 batches, with all batches containing samples from both healthy controls and transplanted animals. The “ComBat” algorithm was implemented to adjust for batch effects (*56*) and probe-sets containing low signal-to-noise measurement were filtered out in order to enhance statistical testing power (*57*). Probe-sets were annotated using 1) annotation file from Dr. Robert B. Norgren Jr (*58*); 2) the annotation file provided by the chip manufacturer (release 33); and 3) data provided by Ingenuity Systems (Ingenuity Systems, www.ingenuity.com) for the small number of probe-sets that were not annotated by the chip manufacturer. Principal Component Analysis (PCA) was applied to summarize gene array variance using the Bioconductor (*59*) MADE4 package (*60*). Analysis of gene differential expression (DE) was performed using an empirical Bayes moderated t-statistic, with a cutoff of 0.05, corrected for multiple hypothesis testing using Benjamini-Hochberg procedure and an absolute fold change cutoff >1.4 with the limma package (*61*).

### Gene Set Enrichment Analysis (GSEA)

GSEA was performed using GSEA 4.1.0 tool (*62, 63*) and gene sets from the Molecular Signatures Database v7.4 both on aggregate sample data from each cohort as a whole. In the current analysis, gene sets were ranked using a signal to noise ratio difference metric with 1000 permutations of gene set labels.

### Functional annotation of genes and pathway enrichment analysis using Metascape

Functional annotation and pathway enrichment analysis was performed using the Metascape web tool (metascape.org) (*41*). First, rhesus macaque genes were mapped to the human orthologs. Then, all statistically enriched terms from gene ontology (GO), KEGG, Biocarta, Reactome, HALLMARK, CORUM, canonical pathways, WikiPathways databases, accumulative hypergeometric p-values and enrichment factors were calculated and used for filtering. Remaining significant terms were then hierarchically clustered into a tree based on Kappa-statistical similarities among their gene memberships. Then 0.3 kappa score was applied as the threshold to cast the tree into term clusters. The terms within each cluster are exported in the Excel spreadsheet named “Enrichment Analysis” (Table S5-S6).

We then selected a subset of representative terms from this cluster and converted them into a network layout. More specifically, each term is represented by a circle node, where its size is proportional to the number of input genes falling into that term, and its color represents its cluster identity (i.e., nodes of the same color belong to the same cluster). Terms with a similarity score >0.3 are linked by an edge (the thickness of the edge represents the similarity score). The network is visualized with Cytoscape (v3.1.2) (*64*) with “force-directed” layout and with edge bundled for clarity. One term from each cluster is selected to have its term description shown as label.

### Analysis of upstream regulators using Ingenuity Pathway Analysis

Pathway analysis on the differentially expressed genes was performed using Ingenuity Pathway Analysis (IPA; https://.qiagenbioinformatics.com/products/ingenuity-pathway-analysis/).

Upstream regulators (limited to the following terms: complex, cytokine, group, growth factor, G-coupled receptor, kinase, ligand-dependent nuclear receptor, micro-RNA, mature micro-RNA, phosphatase, transmembrane receptor, transcriptional regulator, translational regulator) were considered activated or inhibited if enrichment z-scores were greater than 1.25 or less than -1.25, respectively, and p values were <0.05 using the Benjamini-Hochberg-corrected t-test.

### Immunofluorescence analysis of NHP colon

Colon specimens were collected at terminal analysis from different experimental cohorts. Tissues were fixed in 10% buffered formalin and embedded in paraffin. Tissue sections (4 µm) were obtained using a microtome, deparaffinized and then antigen retrieval was performed at 120°C for 30 sec in Trilogy (Cellmarque). Slides were blocked using normal goat serum in PBS containing 0.2% BSA for 1 hour and stained for 1h at 37°C using primary antibodies: rat anti- CD3 (Biorad, Cat# MCA1477) and mouse anti-Ki67 (Dako, Cat# M724029-2). The following fluorescently labeled secondary antibodies were used to stain tissues for 1 hour at 37°C: goat anti-rat AF488 (Invitrogen, Cat# A-11006) and goat anti-mouse IgG1 AF647 (Invitrogen, Cat# A-21240). Sections were washed, stained with Hoechst at 2 µg/mL (Invitrogen, Cat# H3570) and mounted using Prolong Gold (Invitrogen). Images were acquired using a Zeiss Axio Scan Z1 microscope and tile stitching was performed using the ZEN Blue software (Carl Zeiss). Image processing, modeling and quantification of cell populations was performed with the Imaris software v9.7 (Bitplane AG). Among all the tile scan images collected initially (HD= 8, NoRx= 7, REGN421= 9), 5 animals per group presented staining of good quality, with reliable signal for all antibodies and absence of autofluorescence. From each of these tile scan images, 2-6 representative areas per animal were cropped, and 1-3 representative images with well-preserved tissue structure were selected per animal for analysis (total 33 images, with 11 images per group). These images were deidentified and quantification analysis was performed on Imaris in a blinded fashion by a single investigator (D.G.A.). Each channel was processed separately, and cells were modeled as spots based on Hoechst nucleic staining. Data were plotted using Prism (GraphPad) and statistically significant differences determined using Kruskal-Wallis multiple comparison test.

### Mice

C57BL/6 (B6, H-2^b/b^, Thy1.2^+^) and C57BL/6-Thy1.1 mice (B6-Thy1.1, H-2^b/b^, Thy1.1^+^) were originally from The Jackson Laboratory and bred at the University of Pennsylvania. C57BL/6 x BALB/c F1 (model #100007, CBF1, H-2^b/d^, Thy1.2^+^) and congenic B6.SJL-Ptprc^a^ mice (model #002014, B6-CD45.1, H-2^b/b^, Thy1.2^+^) were from The Jackson Laboratory (Bar Harbor, ME). B6 *Cd4-Cre*;*ROSA^DNMAML^* mice (expressing the pan-Notch inhibitor DNMAML fused to GFP in mature CD4^+^ and CD8^+^ T cells under the control of the *ROSA26* promoter) were described previously (*19, 65*). *Itgb1^f/f^* mice on a mixed B6;129 background (H-2^b/b^) were from The Jackson Laboratory (model #004605) (*66*). *Itgb1^f/f^* mice were crossed to *Cd4-Cre*;*ROSA^DNMAML^* mice to generate *Cd4-Cre;Itgb1^f/f^* vs. *Cd4-Cre;Itgb1^f/f^;ROSA^DNMAML^* mice. For bone marrow transplantation experiments, co-housed littermate controls were used as donors and recipients at the University of Pennsylvania, per protocols approved by the University of Pennsylvania’s Office of Regulatory Affairs.

### Mouse study design and transplantation strategy

For transplantation, CBF1 recipient mice were irradiated using a Cesium-137 source (11 Gy, split into two 5.5 Gy doses separated by 3h). Recipients were 8-16 weeks old. Both female and male mice were used as recipients and equally distributed among experimental groups. T cell-depleted BM (TCD BM) prepared with anti-Thy1.2 antibodies and complement (Cedar Lane Laboratories) was injected i.v. with or without wild type or DNMAML T cells purified by EasySep negative magnetic selection (Stem Cell Technologies), as described (*21*). In selected experiments, suspensions of splenocytes and lymph node cells were transplanted without further T cell purification.

### Mouse tissue processing

Single cell suspensions from spleen, mesenteric lymph nodes, or pooled peripheral lymph nodes (cervical, brachial, axial, inguinal) were prepared by physical disruption through 70 μm cell strainers. Single cell suspensions were prepared from liver as previously described (*17*). Intestinal lamina propria and intestinal epithelium was prepared as previously described with slight modifications (*19*). Briefly, intestines were washed and cut longitudinally after removal of Peyer’s patches from the small intestine. Fragments (0.5-1 cm) of intestine were cleaned of mucus and feces and incubated in phosphate-buffered saline with 2mM EDTA and 5% FBS without dithiothreitol for 45′ while shaking at 37°C. Supernatant was passed through 100μM MACS SmartStrainers (Miltenyi) and collected as intestinal epithelial lymphocyte (IEL) fraction. For isolation of lamina propria lymphocytes (LPL), the remaining tissue was incubated in RPMI/FBS 10% with 1mg/mL collagenase IV (Gibco) for 20′ while shaking at 37°C. The resulting tissue was vigorously pipetted and supernatant passed through 100μM MACS SmartStrainers. Debris was removed from liver and intestinal fractions using 40%:80% Percoll gradient separation.

### Multiparameter flow cytometry on mouse samples

The following antibodies were from Biolegend: anti-CD4 (clone GK1.5); CD8α (clone 53-6.7); CD44 (clone IM7); CD45.1 (clone A20); CD45.2 (clone 104); Thy1.1 (clone OX-7); Thy1.2 (clone 30-H12); GFP (AF488-labeled cloneFM264G); α4β7 (LPAM-1, clone DATK32); integrin α4 (clone R1-2); integrin β1 (clone HMβ1-1); integrin β7 (clone FIB27). Anti-CD25 (clone PC61) was from BD Biosciences. Anti-Foxp3 (clone FJK-16s) was from eBioscience. Fc receptors were blocked with unlabeled anti-mouse CD16/32 (BioLegend). Non-viable cells were excluded from analysis with Zombie Aqua Fixable Viability Dye (Biolegend), or DAPI (Sigma Aldrich). For intranuclear staining, Foxp3/Transcription Factor Staining Buffer (eBioscience) was used per manufacturer’s protocol. Flow cytometric analysis was performed using a 4 or 5- laser BD Fortessa (Becton Dickinson). Analysis was performed using FlowJo software (Becton Dickinson).

### Statistical analysis

All data was analyzed using Prism 9 (GraphPad). Survival data were analyzed using log-rank (Mantel-Cox) test. Flow cytometry data, immunofluorescent imaging data, and histological data were analyzed using one-way or two-way ANOVA with Tukey post-hoc multiple comparison test, or using Kruskal-Wallis tests, there appropriate. Technical and biological replicates are indicated in the figure legends. Error bars on the plots represent standard error of the mean (SEM). p < 0.05 was considered as significant. Details of statistical methods used for each analysis are provided in the corresponding figure legends. Details of the statistical analysis of the gene array data are provided in the corresponding section.

## Supplementary Materials

**Fig. S1**. Pharmacokinetics of REGN421 in the allo-HCT NHP model and hematopoietic reconstitution of REGN421-prophylaxed allo-HCT recipients.

**Fig. S2**. T cell reconstitution in the REGN421 experimental cohort.

**Fig. S3**. Expression of chemokine receptors on CD8^+^ and CD4^+^ T cells.

**Fig. S4**. Cell-intrinsic canonical Notch signals control α4β7 expression and gut-homing potential in alloreactive T cells.

**Fig. S5**. Cell surface individual integrin subunits in NHP splenic T cells after allo-HCT.

**Fig. S6**. Loss of integrin β1 rescues α4β7 expression in Notch-deprived alloreactive T cells.

**Fig. S7**. Working model.

**Table S1**. Transplant characteristics and data inclusion.

**Table S2**. List of DE genes.

**Table S3**. aDLL4 specific DE genes.

**Table S4**. aDLL4-aOX40L-aCD28 shared DE genes

**Table S5**. Shared signature pathway enrichment analysis

**Table S6**. aDLL4 signature pathway enrichment analysis

**Table S7**. aDLL4 signature upstream regulators.

**Table S8**. List of critical reagents.

## Acknowledgments

### Funding

This work was supported by the following sources: Translational Research Program grant from the Leukemia and Lymphoma Society TRP-6583-20 (L.S.K. and I.M.)

National Institutes of Health R01-HL095791 (L.S.K.)

National Institutes of Health P01-HL158504 (L.S.K.)

National Institutes of Health U19-AI051731 (L.S.K.)

National Institutes of Health R01-AI091627 (I.M.)

National Institutes of Health R37-AI34495 (B.R.B.)

National Institutes of Health R01-HL56067 (B.R.B.)

National Institutes of Health R01-HL11879 (B.R.B.)

National Institutes of Health R01-HL-115114 (B.R.B.)

Research funding from Regeneron, Inc.

Be the Match Foundation/CIBMTR Amy Strelzer Manasevit Research Grant (V.T.)

ASTCT New Investigator Award (V.T.),

National Institutes of Health T32-GM007863 (E.P.)

National Institutes of Health T32-AI070077 (A.V.)

National Institutes of Health F30-AI161873 (A.V.)

National Institutes of Health F30-AI136325 (E.P.).

### Author contributions

V.T., A.V. and E.P. designed all research studies, performed experiments and analyzed data. C.M., D.G.A, U.G., X.R., J.L., D.J.H., H.Z., M.H., A.Y., S.K., A.A. and G.C. performed experiments and analyzed data. V.T. and S.F. performed bioinformatic analysis. A.P-M. performed histopathological scoring. B.B., Y.S. provided critical reagents and advice. S.M.C., O.H., F.K., G.T. and B.R.B. provided critical reagents and/or advice with experimental design and data interpretation. L.S.K. and I.M. selected the research questions, defined the overall experimental approach and supervised the work. V.T., L.S.K. and I.M. wrote the manuscript, and all authors reviewed it.

### Data and materials availability

Rhesus macaques microarray data are deposited in GEO (GSE…). All other data are available in the main text or the supplementary materials.

---

**Figure S1.**
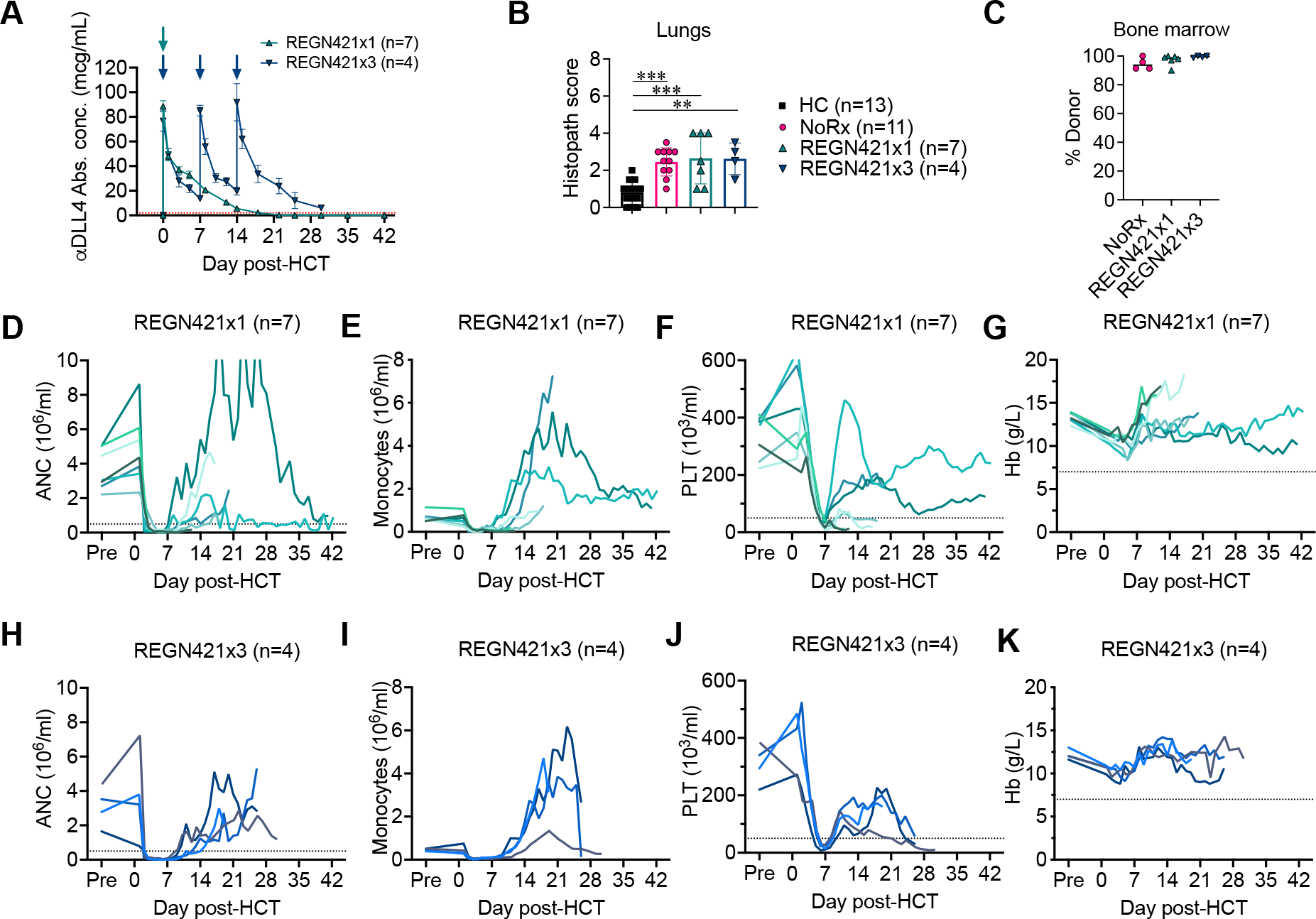
Pharmacokinetics of REGN421 in the allo-HCT NHP model and hematopoietic reconstitution of REGN421-prophylaxed allo-HCT recipients. (A) Systemic concentrations of REGN421 following the single or triple dosing regimen (REGN421x1, REGN421x3). (B) Histopathological aGVHD scores for lungs. **p<0.01, ***p<0.001 using one-way ANOVA with Tukey post-hoc-test. (C) Donor chimerism in the bone marrow at the time of terminal analysis. (D-K) Absolute numbers of neutrophils (ANC; plots D and H), monocytes (plots E and I), platelets (PLT, plots F and J) and hemoglobin (Hb) levels (G and K) in REGN421x1 (plots D-G) and REGN421x3 (plots H-K) cohorts.

**Figure S2.**
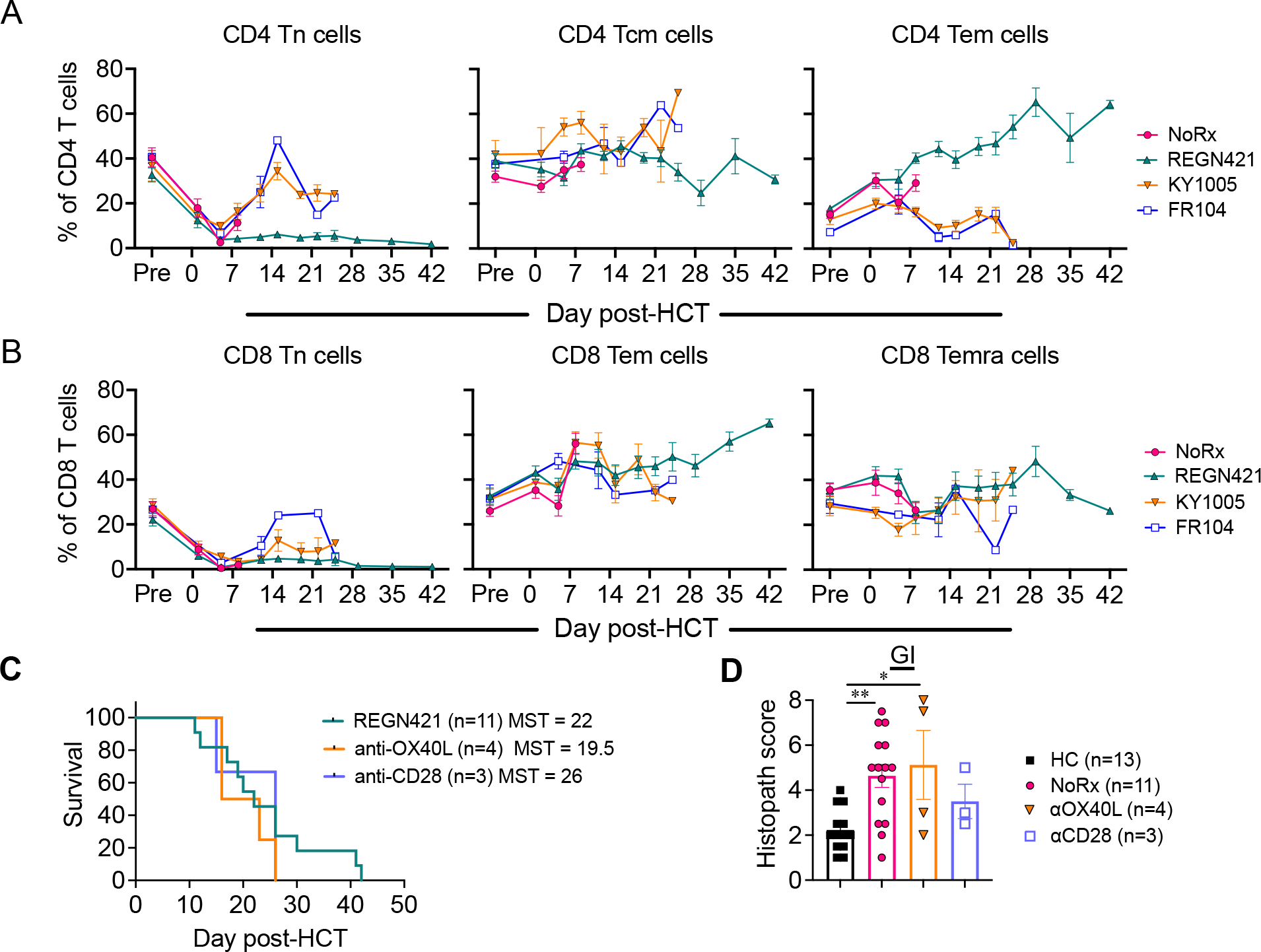
T cell reconstitution in the REGN421 experimental cohort. (A-B) Absolute numbers of CD4^+^ (A) and CD8^+^ (B) T cells with CD45RA^+^CCR7^+^CD95^−^ naïve, CD45RA^−^ CCR7^+^ central memory and CD45RA^−^CCR7^−^ effector-memory phenotypes in the peripheral blood of allo-HCT recipients. (C) Overall survival of allo-HCT recipients in the combined REGN421 cohort (recipients that received either one or three doses of REGN421, n=11), anti- CD28 (purple, n=3) and anti-OX40L experimental cohort (orange, n=4). (D) Histopathological aGVHD scores for GI tract in the combined REGN421 cohort (recipients that received either one or three doses of REGN421, n=11), anti-CD28 (purple, n=3), and anti-OX40L experimental cohort (orange, n=4) **p<0.01, ***p<0.001 using one-way ANOVA with Tukey post-hoc-test.

**Figure S3.**
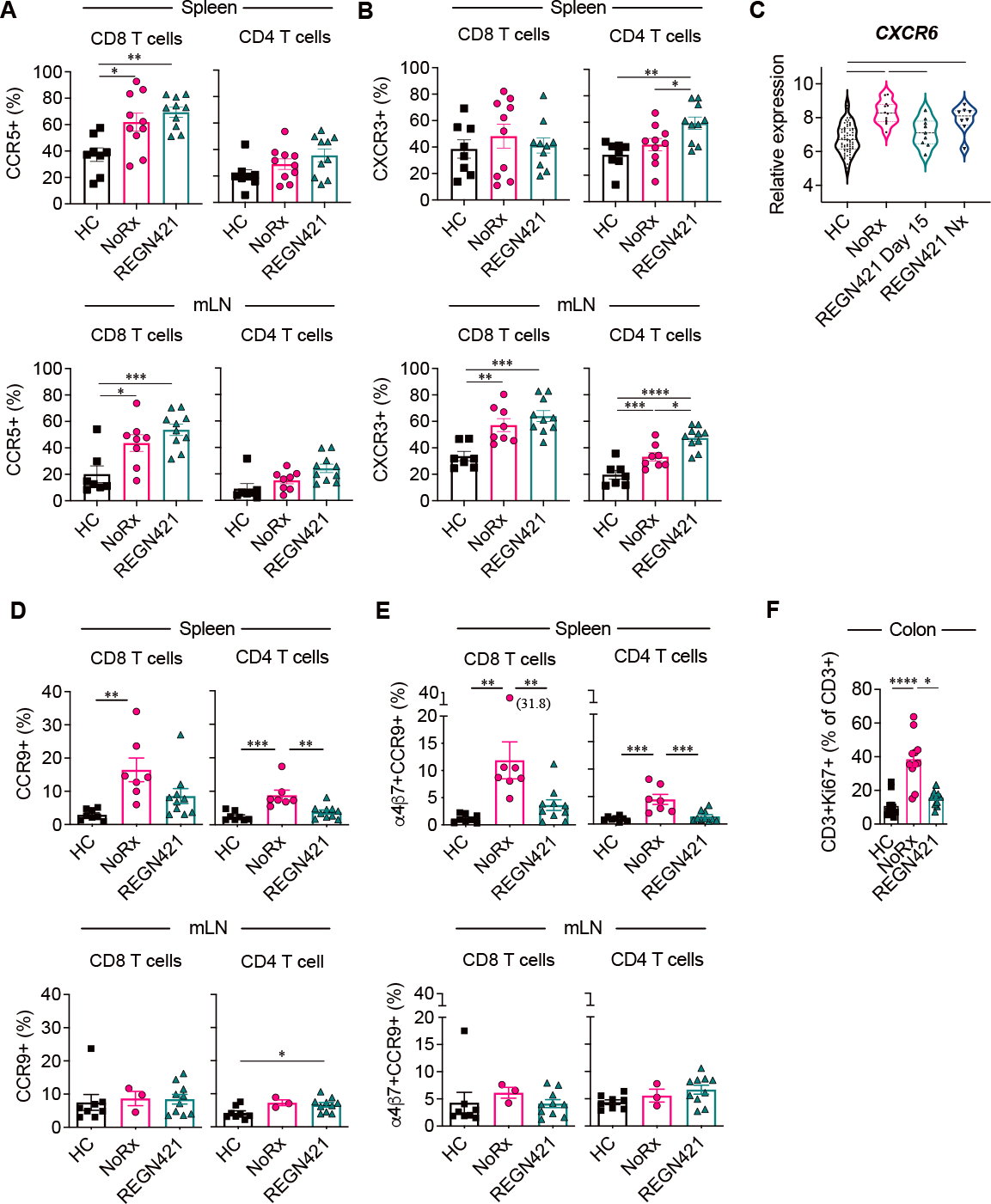
Expression of chemokine receptors on CD8^+^ and CD4^+^ T cells. (A-B) Relative number of CCR5^+^ (A) or CXCR3^+^ (B) CD8^+^ and CD4^+^ T cells in the spleen and mesenteric lymph nodes of healthy control animals (n=7-8 depending on organ) in allo-HCT recipients from NoRx aGVHD (n=8-10 depending on the organ) vs. REGN421 (n=10) cohorts. (C) Relative abundance of *CXCR6* mRNA in CD3^+^CD20^−^ T cells from peripheral blood, sorted on day +15 after allo-HCT or at the time of terminal analysis. (D-E) Relative number of CCR9^+^ (D) or a4b7^+^CCR9^+^ (E) CD8^+^ and CD4^+^ T cells in the spleen and mesenteric lymph nodes of healthy control animals (n=8 depending on organ) in allo-HCT recipients from NoRx aGVHD (n=3-7 depending on organ) vs. REGN421 (n=10) cohorts. *p<0.05,**p<0.01,***p<0.001 using one-way ANOVA with Tukey post-hoc-test. (F) Proportion of Ki67^+^ cells among CD3^+^ T cells in the colon as assessed by immunofluorescence imaging and quantified using the Imaris software (n=11 images from 5 animals per group). Kruskal-Wallis multiple comparison test with *p<0.05, ****p<0.0001.

**Figure S4.**
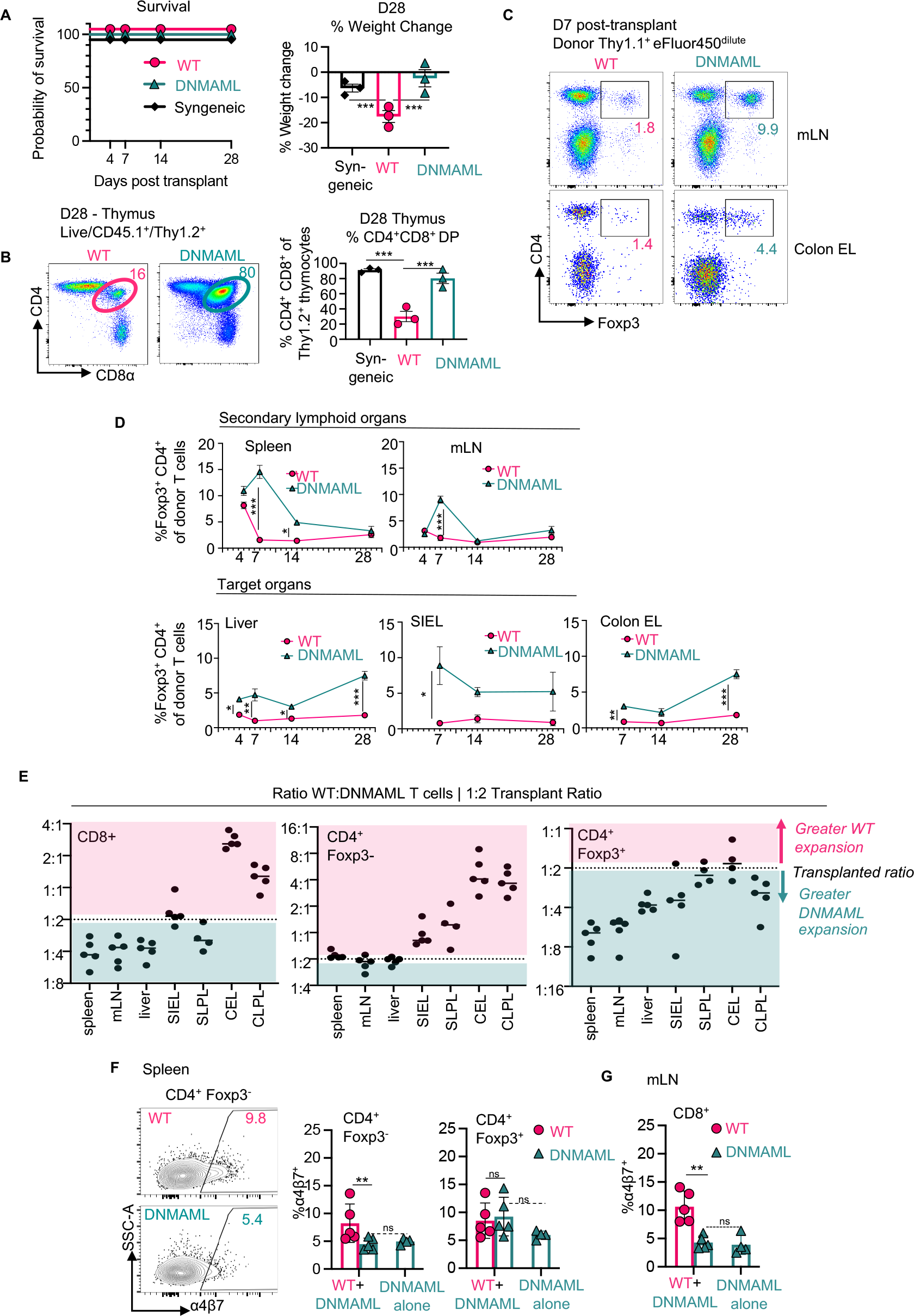
Cell-intrinsic canonical Notch signals control α4β7 expression and gut-homing potential in alloreactive T cells. (A-D) These data are related to the experiments depicted in Fig. 5A-C. (A) Survival and weight change at day 28 post-transplant. (B) Representative flow cytometry plots showing the frequency of CD4^+^CD8^+^ cells among newly formed CD45.1^+^Thy1.2^+^ thymocytes at day 28 post-transplant, summarized on the right (as readout for thymic GVHD). (C) Flow cytometry plots showing the frequency of Foxp3^+^CD4^+^ Tregs among donor T cells in mLN and colon epithelium, as summarized in multiple organs across multiple timepoints in (D). (E) Similar experiment as in Fig. 5F, but recipients were transplanted with a 1:2 ratio of wild type:DNMAML T cells. Relative accumulation of wild type to DNMAML T cells was calculated in each organ fraction. (F) Flow cytometric analysis of α4β7 expression in donor Foxp3^-^CD4^+^ conventional T cells from recipients receiving wild type and DNMAML T cells (1:1) vs. DNMAML T cells alone in spleen (F) and mLN (G). *p<0.05, **p<0.01, ***p<0.001, one-way ANOVA with Tukey’s post-hoc-tests. Syn = syngeneic, mLN = mesenteric lymph node, SI = small intestine, C = colon, EL = epithelial lymphocytes, LPL = lamina propria lymphocytes.

**Figure S5.**
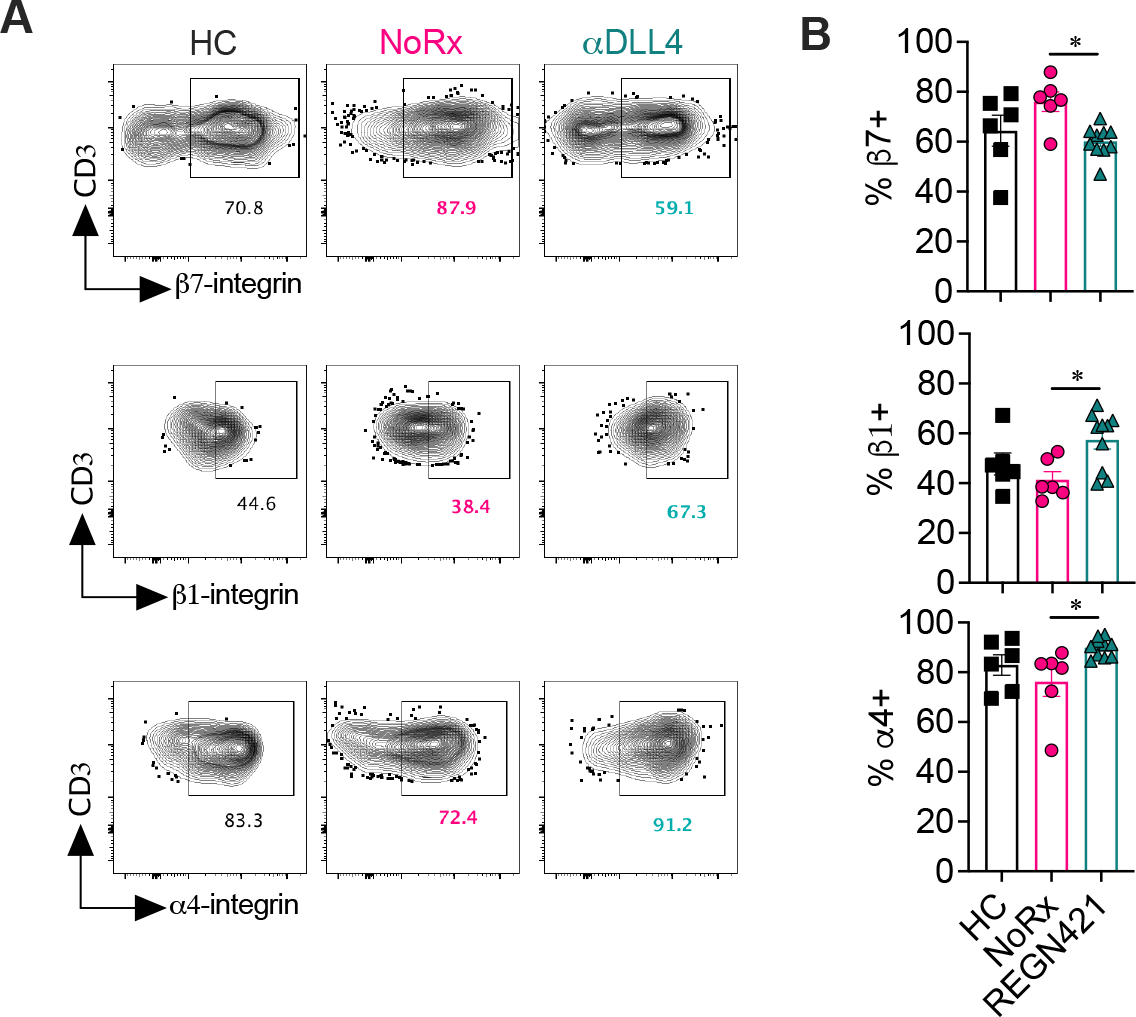
Cell surface individual integrin subunits in NHP splenic T cells after allo-HCT. Flow cytometric analysis of α4, β1 and β7-integrin subunit expression among spleen CD8^+^ T cells at day 15 after transplantation in allo-HCT recipients as compared to healthy controls (HC). Findings are shown for unprophylaxed aGVHD (NoRx, n=6) and REGN421-treated (n=10) cohorts, with representative flow cytometry plots (A) and resulting data (B). *p<0.05 using one-way ANOVA with Tukey’s post-hoc-tests.

**Figure S6.**
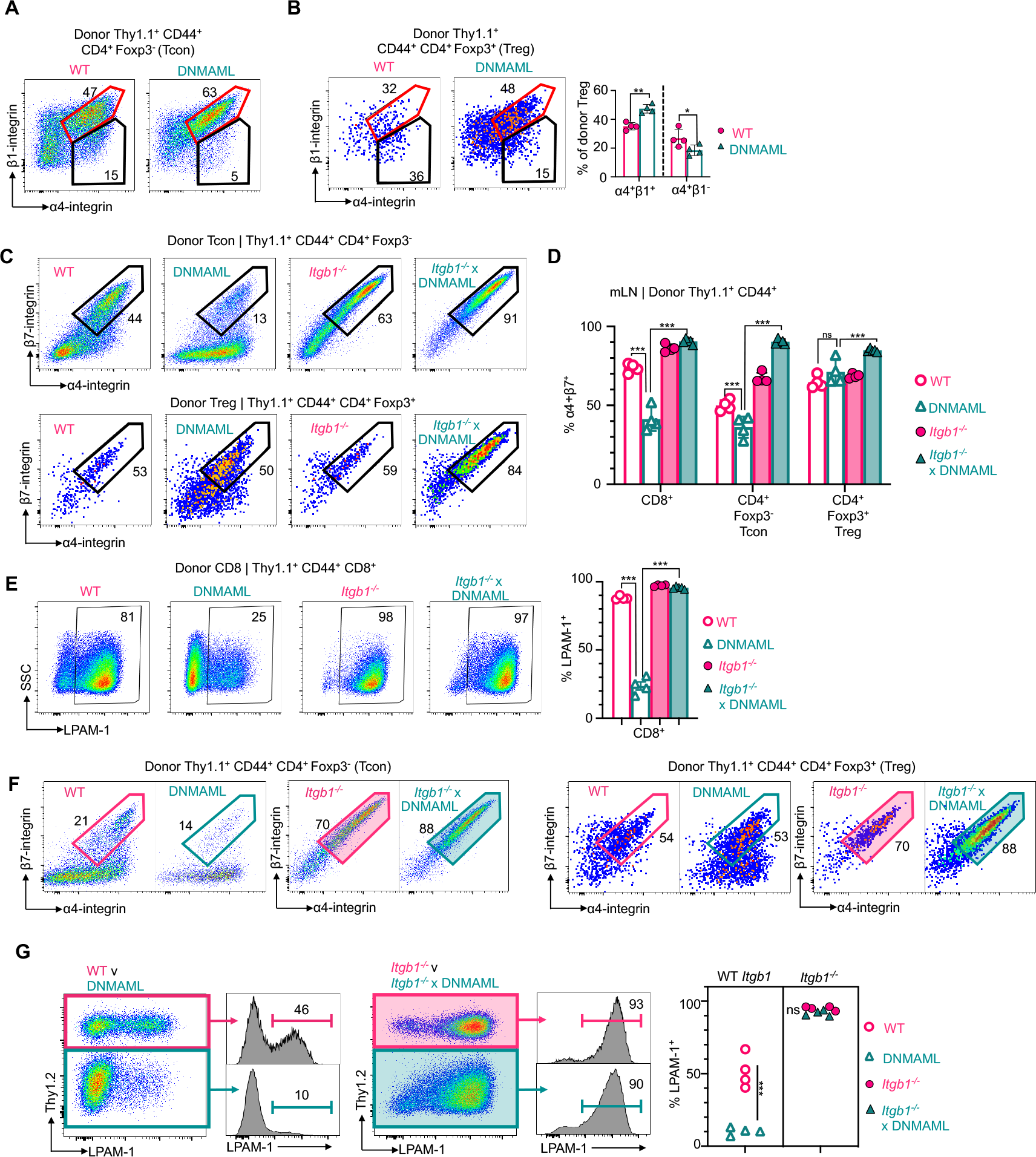
Loss of integrin β1 rescues α4β7 expression in Notch-deprived alloreactive T cells. (A-E) These data are related to the experiment depicted in Fig. 7A–E. Representative flow plots showing α4β1 expression among donor alloreactive CD44^+^CD4^+^ Foxp3^-^ (Tcon) (A) and Foxp3^+^ (Treg) (B). (C) Representative flow cytometry plots showing α4β7 expression in CD4^+^ Tcon (top) and Treg (bottom) represented as summary data in Fig. 7E. (D) Summary data of α4β7 expression in donor T cell subsets harvested from mesenteric lymph nodes. (E) Representative flow cytometry plots and summary data for a4b7/LPAM-1 heterodimer expression in donor CD8^+^ cells. (F-G) Related to experiment from Fig 7F-G. (F) Representative flow cytometry plots of cell surface α4β7 in competing donor CD4^+^ Tcon (left) and Treg (right) subsets as summarized in Fig. 7G. (G) Representative flow cytometry plots and summary data of α4β7/LPAM-1 abundance in competing donor CD8^+^ cells and associated summary data (right). n=4 mice per group for both experiments., ***p<0.001, two-way ANOVA with Tukey’s post-hoc-tests. ns = not significant. mLN = mesenteric lymph node.

**Figure S7.**
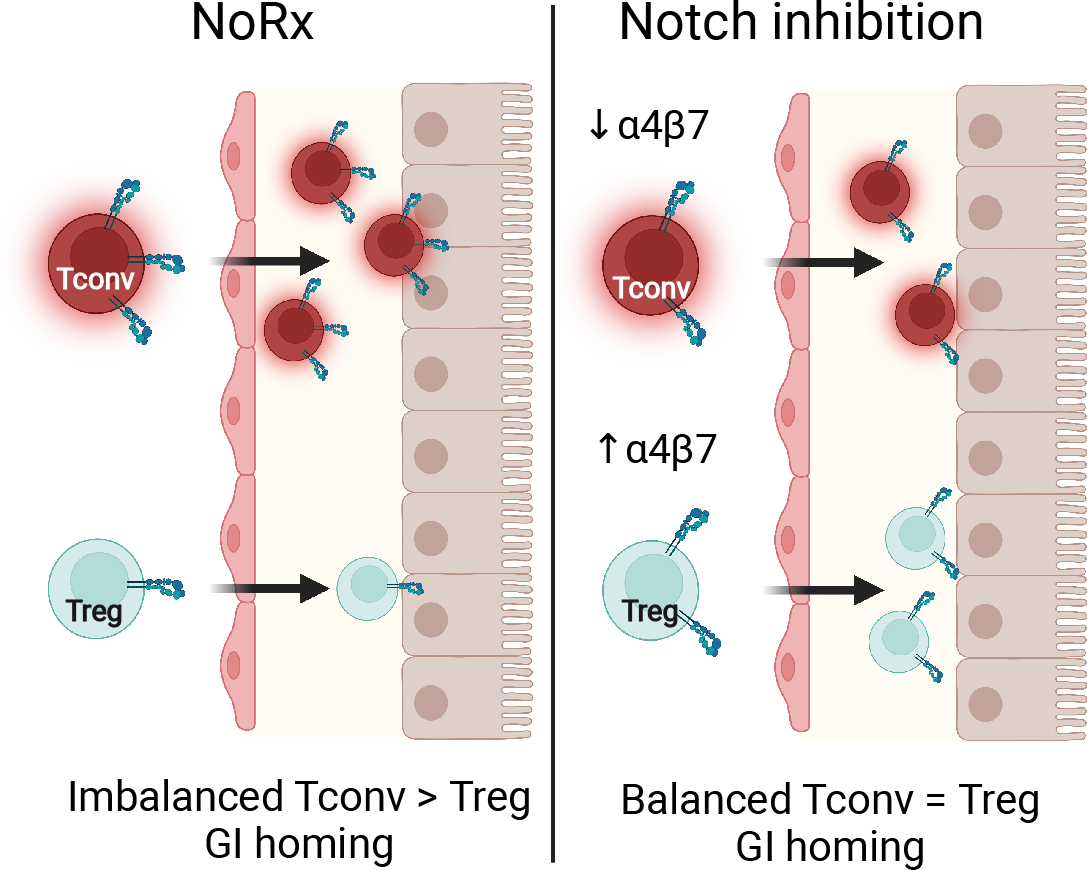
Working model depicting how inhibition of Notch signaling in T cells differentially affects cell surface α4β7/LPAM-1 in Tcon and Treg, resulting in decreased Tcon gut infiltration and an increased gut Treg/Tcon ratio early after allo-HCT.

